# Sleepiness, not total sleep amount, increases seizure risk

**DOI:** 10.1101/2023.09.30.560325

**Authors:** Vishnu Anand Cuddapah, Cynthia T Hsu, Yongjun Li, Hrishit M Shah, Christopher Saul, Samantha Killiany, Joy Shon, Zhifeng Yue, Gabrielle Gionet, Mary E Putt, Amita Sehgal

**Affiliations:** Epilepsy Neurogenetics Initiative, Division of Neurology, Departments of Pediatrics and Neurology, Children’s Hospital of Philadelphia; Philadelphia, PA 19104; Department of Neurology, Perelman School of Medicine, University of Pennsylvania; Philadelphia, PA 19104; Howard Hughes Medical Institute, University of Pennsylvania; Philadelphia, PA 19104; Chronobiology and Sleep Institute, Perelman School of Medicine, University of Pennsylvania; Philadelphia, PA 19104; Department of Biostatistics, Epidemiology and Informatics University of Pennsylvania; Philadelphia, PA 19104; CHOP/Penn Intellectual and Developmental Disabilities Research Center; Philadelphia, PA 19104

**Author notes:** Corresponding author: Amita Sehgal.

## Abstract

Sleep loss has been associated with increased seizure risk since antiquity. Despite this observation standing the test of time, how poor sleep drives susceptibility to seizures remains unclear. To identify underlying mechanisms, we restricted sleep in *Drosophila* epilepsy models and developed a method to identify spontaneous seizures using quantitative video tracking. Here we find that sleep loss exacerbates seizures but only when flies experience increased sleep need, or *sleepiness*, and not necessarily with reduced sleep quantity. This is supported by the paradoxical finding that acute activation of sleep-promoting circuits worsens seizures, because it increases sleep need without changing sleep amount. Sleep-promoting circuits become hyperactive after sleep loss and are associated with increased whole-brain activity. During sleep restriction, optogenetic inhibition of sleep-promoting circuits to reduce sleepiness protects against seizures. Downregulation of the 5HT1A serotonin receptor in sleep-promoting cells mediates the effect of sleep need on seizures, and we identify an FDA-approved 5HT1A agonist to mitigate seizures. Our findings demonstrate that while homeostatic sleep is needed to recoup lost sleep, it comes at the cost of increasing seizure susceptibility. We provide an unexpected perspective on interactions between sleep and seizures, and surprisingly implicate sleep-promoting circuits as a therapeutic target for seizure control.

## Introduction

Since the writings of Hippocrates and Aristotle into the modern era, poor sleep has been associated with increased seizure risk^1^. People with epilepsy, as well as caregivers of children with epilepsy, report that sleep disruption is a common trigger for seizure exacerbation^2, 3^. Understanding how sleep restriction might increase seizure risk has been challenging because of multiple confounding variables that change through the day, including light exposure, food intake, sleep and wake, physiological processes regulated by endogenous circadian clocks, etc. However, when well-defined protocols are used, distinct contributions of sleep on brain excitability have been suggested^4, 5^.

To understand how sleep restriction increases seizure severity, we leveraged the tractability of *Drosophila*. Sleep loss leads to diffuse and focal effects in the *Drosophila* brain. On a broad level, synapse number and size increase in the fly brain with increased time awake^6^. At a circuit/cellular level, sleep deprivation leads to increased activity of the dorsal fan-shaped body (dFB)^7, 8^ and ellipsoid body^9^, which drives increased sleep. Specific Kenyon cells in the mushroom body (MB)^10–12^, subsets of neurons in the pars intercerebralis (PI)^13, 14^, and 2 cholinergic neurons in the ventral nerve cord (VNC)^15, 16^ also have sleep-promoting roles. In addition, several loci in the fly brain drive wakefulness^8, 12, 17^. In the present study, we investigated if activity of sleep:wake-regulating circuits modulates seizure risk.

### Sleep restriction worsens induced seizures

To first understand if sleep loss is associated with seizure severity in *Drosophila*, we took advantage of bang-sensitive neurogenetic models of epilepsy. Bang-sensitive flies are prone to having seizures after sustaining a strong mechanical stimulus, e.g. after flies are banged against a countertop or vortexed. Like mammalian models, seizures in flies occur as a sequence of stereotyped behaviors: 1) an atonic “paralytic” phase, 2) a tonic/clonic “convulsive” phase, and a 3) post-ictal recovery phase (Ext. Data Fig. 1a)^18^. While most studies monitor the total time to recovery from seizures, we sought to assess seizure severity by timing how long each fly spent in each of these discrete phases; this was achieved through video recording followed by time measurements (Ext. Data Fig. 1a and Supplementary Video 1). *tko*^25t^ mutant flies exhibit bang-sensitive seizures and were sleep restricted through caffeine feeding as previously described^19^. Caffeine led to sleep loss, and this was associated with prolonged seizures and recovery times in *tko*^25t^ flies (Ext. Data Fig. 1b-e, Ext. Data Fig. 2a-d). We also assessed *eas^pc^*^80f^ bang-sensitive flies, which demonstrated a loss of nighttime sleep after caffeine exposure and an increase in total seizure time (Ext. Data Fig. 1f-i, Ext. Data Fig 2e-h). In the absence of caffeine-induced sleep loss, *eas^pc^*^80f^ mutant flies did not exhibit a tonic/clonic phase; this phase only appears after sleep restriction (Ext. Data Fig. 2n). Caffeine treatment led to increased fly death in bang-sensitive mutants but not wild-type flies (Ext. Data Fig. 2j, l, o), correlating with increased seizure severity. These data demonstrate that sleep restriction worsens seizures in flies consistent with previous findings^20^.

We considered whether other methods of sleep restriction also led to worsened seizures. We initially used mechanical stimulation to restrict sleep, but this resulted in bang-sensitive seizures during the protocol, pushing flies into a subsequent refractory state resistant to seizure induction. We then turned to genetic manipulations to restrict sleep. *tko*^25t^ mutant flies were crossed to *redeye* mutant flies (*rye*) or *sleepless* mutant flies (*sss^p^*^1^), which are short-sleeping mutant flies exhibiting a significant reduction in sleep amount. *tko*^25t^ flies crossed to these short-sleeping mutants exhibited a decrease in sleep with prolongation of seizure times, mostly driven by a longer tonic-clonic phase (Ext. Data Fig. 1j-q; Ext. Data Fig. 3a-j). Taken together, these data demonstrate that both pharmacologically- and genetically-induced sleep loss leads to worsened seizures.

Given that sleep loss is associated with seizure exacerbation, we hypothesized that sleep enhancement would protect against seizures. To test this hypothesis, we fed bang-sensitive flies gaboxadol (also known as THIP), a GABA-A receptor agonist known to promote sleep^21^, and measured seizure severity. Gaboxadol treatment reliably increased sleep amount across genotypes (Ext. Data Fig. 1b-d, f-h); surprisingly, it was not protective against seizures and instead prolonged seizures in *eas^pc^*^80f^ mutant flies (Ext. Data Fig. 1e, i). Given likely off-target effects of gaboxadol, we also sought to enhance sleep using a more specific genetic approach. Overexpression of UAS-*nemuri* using an inducible, pan-neuronal Gal4 driver (*elav*-GeneSwitch, *elav*-GS) leads to increased sleep^22^. Induction of neuronal UAS-*nemuri* expression in the *tko*^25t^ background by feeding flies the GeneSwitch activator RU486 led to an increase in sleep (Ext. Data Fig. 4a-c). This sleep induction led to a complete block of seizures (Ext. Data Fig. 4d). A smaller sleep induction was also noted in the absence of RU486 (Ext. Data Fig. 4b), likely indicating leaky *nemuri* expression. Even this small sleep induction was sufficient to decrease seizure times in *tko*^25t^ flies (Ext. Data Fig. 4e-i). To determine if the protection against seizures was due to a direct effect of *nemuri* or an effect of increased sleep, *tko*^25t^ flies were crossed with *nemuri* mutant flies (*nur*^3^). Sleep was similar in *tko*^25t^ and *tko*^25t^*/nur*^3^ flies (Ext. Data Fig. 5a-c), consistent with previous findings that loss of *nemuri* does not change daily sleep amount^22^. Loss of *nemuri* did not have a significant effect on seizures (Ext. Data Fig. 5d-i). This is consistent with the conclusion that sleep enhancement, and not direct effects of *nemuri*, is responsible for seizure protection. Overall, while sleep enhancement with gaboxadol promoted seizures, sleep enhancement with *nemuri* blocked seizures. This indicated that other variables related to sleep beyond purely sleep duration might be important drivers of seizure severity.

### Sleep loss leads to spontaneous seizures

Unexpectedly, while caffeine-induced sleep loss increased seizure severity, it decreased the likelihood of induced seizures occurring (Ext. Data Fig. 2m), similar to previously noted effects of mechanical sleep restriction on temperature-sensitive seizures^23^. Bang-sensitive seizures are known to be less likely if flies are in a refractory state due to recent seizures^24, 25^; this raised the possibility that undocumented spontaneous seizures were occurring prior to the bang-sensitive assay while the flies were being sleep-deprived with caffeine. The occurrence of spontaneous seizures has been suspected in fly models of epilepsy^26^, but the lack of non-invasive electroencephalography has made this challenging to confirm. Therefore, we sought to create a method to identity spontaneous seizures in flies.

We developed a novel chronic video-tracking pipeline to automatically detect and quantify spontaneous, previously undocumented, seizures in flies (Fig. 1a). In this paradigm, flies were placed into individual wells of a 24- or 48-well plates and video recorded for 96 hours in 12-hour light:12-hour dark cycles using an infrared camera at 25 frames per second. Video tracking software was used to convert fly body positions to XY coordinates over time. We then identified velocity, acceleration, and pixel change thresholds that are typical of convulsive and never occur in Canton-S or *w1118* wild-type flies (CynthiSeize algorithm; see Supplementary Methods). This pipeline not only enabled robust identification of spontaneous (non-induced) seizures but also simultaneous quantification of sleep state, sleep amount, and seizure lethality for individual flies (see Supplementary Video 2 for sample of spontaneous seizure). To further validate that the identified hyperkinetic tonic-clonic behavioral sequences were indeed seizures, we pan-neuronally expressed TRIC-luciferase, a GFP-based calcium reporter coupled to luciferase that allows for measurement of neural activity in awake behaving flies^27^. This demonstrated that the hyperkinetic tonic-clonic behavioral sequence in flies was associated with prolonged neuronal hypersynchronous neuronal activation (Ext. Data Fig. 7a-h), consistent with seizures.

**Fig. 1.**
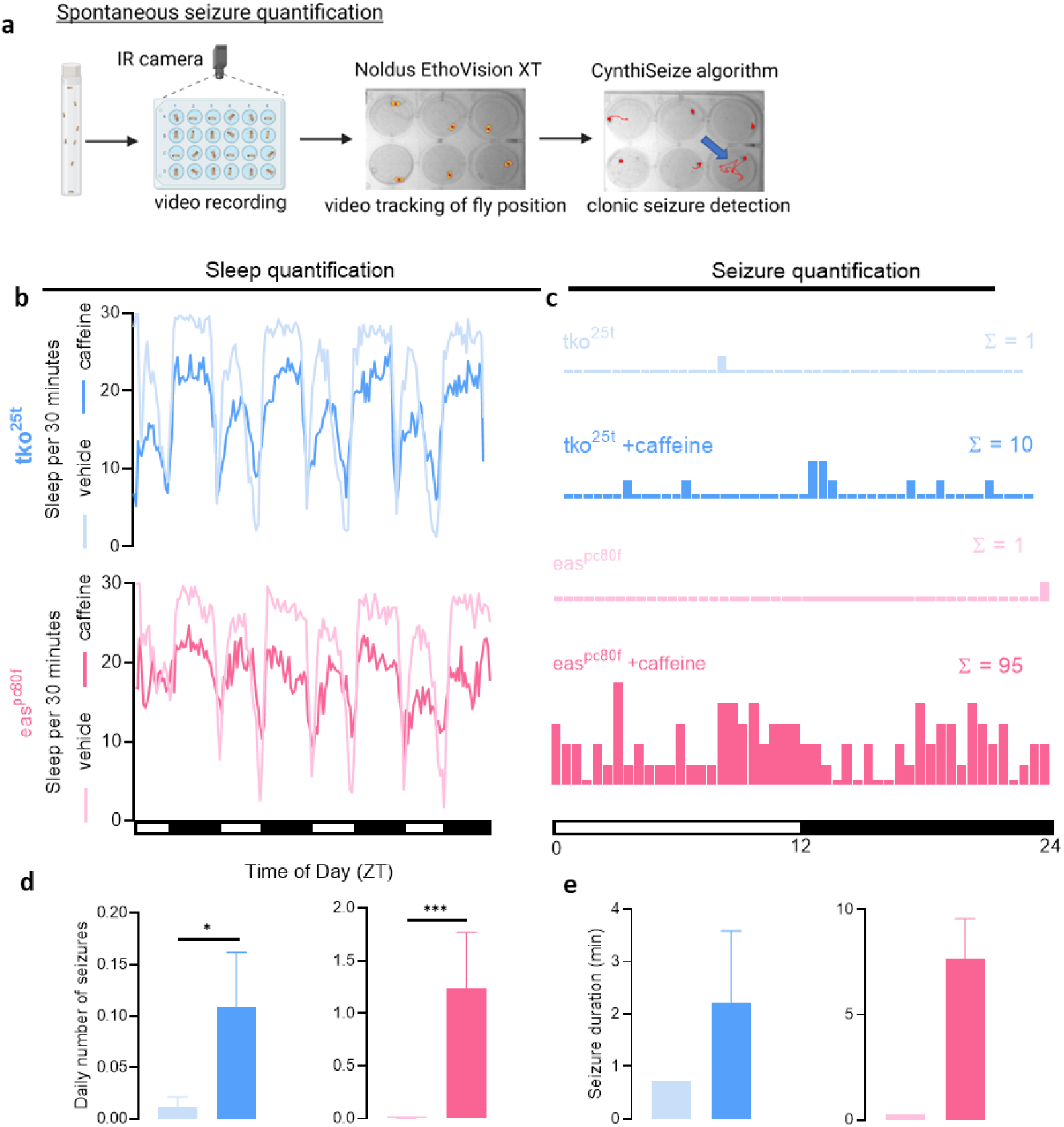
*Drosophila* exhibit spontaneous seizures after sleep restriction. **a**, Flies are load into individual wells of a 24- or 48-well plate then video recorded for 96 hours in a 12-hour light:12- hour dark cycle. EthoVision XT is used to detect fly position (*yellow with red dot)*. An algorithm (“CynthiSeize”) was developed to detect movements consistent with hyperkinetic seizures. Red line tracing = fly position over previous 10 seconds; Blue arrow = fly with sample seizure (*right*). **b**, *tko*^25t^ (*blue*), and *eas^pc^*^80f^ *(pink)*, flies were fed vehicle or caffeine and sleep was measured over multiple day (white bar in *y-axis*) and night (dark bar in *y-axis*) cycles. Caffeine treatment led to less sleep across both models. n=29-32 flies/condition. **c**, Spontaneous seizures were counted across 48 30-minute time bins for each treatment condition in *tko*^25t^ (*blue*) and *eas^pc^*^80f^ *(pink)* flies. Sleep loss with caffeine treatment increased the number of spontaneous seizures (Σ) across both genotypes. n=29-32 flies/condition. **d**, Average number of seizures per day per fly in *tko*^25t^ (*blue*) and *eas^pc^*^80f^ *(pink)* flies is increased by caffeine across genotypes. n=29-32 flies/condition. **e**, Average seizure duration trends upward in *tko*^25t^ (*blue*) and *eas^pc^*^80f^ *(pink)* flies. Statistical testing not possible due to only one fly with a seizure in conditions without caffeine. Error bars not depicted given only one seizure detected in conditions without caffeine. n=1-82 seizures/condition. Negative binomial model with Wald test for daily number of seizures was used. *p<0.05, ***p<0.001. Data are presented as mean values ± SEM.

Using chronic video-tracking, we confirmed that caffeine treatment led to sleep loss in *tko*^25t^ and *eas^pc^*^80f^ mutant flies (Fig. 1b, Ext. Data Fig. 6a, e); in addition, we identified more frequent and severe spontaneous seizures. While spontaneous seizures in bang-sensitive mutants were very rare, sleep restriction with caffeine caused an increase in seizure number and duration across genotypes (Fig. 1c-e). Seizures were more likely to occur when flies were awake (Ext. Data Fig. 6b, f), which correlated with decreased sleep amount in the preceding 3 hours (Ext. Data Fig. 6c, g). Lethal seizures were also more likely to occur when there was decreased sleep in the preceding 3 hours (Ext. Data Fig. 6d, h). These data confirm that bang-sensitive, neurogenetic models of fly epilepsy exhibit spontaneous seizures, and that spontaneous lethal seizures are more likely to occur following prolonged wakefulness.

Beyond neurogenetic forms of epilepsy, we sought to understand if the effects of sleep restriction on seizures were generalizable to other forms of seizures. We fed picrotoxin, a GABA-A receptor antagonist to wild-type Canton-S flies and performed continuous video tracking (see Supplementary Video 3). Even in the absence of a sleep-depriving stimulus, retrospective analyses revealed that seizures were more likely to be lethal when individual flies were awake and had slept less in the preceding three hours (Ext. Data Fig. 8a-c). Seizures were more frequent during wakefulness (Ext. Data Fig. 8d), possibly due to decreased recent sleep (Ext. Data Fig. 8e). Combined with the results from neurogenetic models of seizures, these data indicate that the total amount of sleep achieved and/or the need for sleep may be modulators of seizure severity.

### Sleep need, not total sleep time, controls seizure severity

To disambiguate the effects of sleep history (i.e. how much an organism has slept) versus sleep need (i.e. how much sleep an organism requires) on seizure severity, we took advantage of two thermogenetic models of sleep loss that have variable effects on sleep need. In the *tko*^25t^ background, we drove expression of UAS-TrpA1, a temperature-sensitive cation channel, using the c584-Gal4 and Tdc2-Gal4 drivers. With temperature elevation, c584>TrpA1 leads to sleep loss followed by a homeostatic increase in sleep, indicating that sleep need has accumulated^28^. Specifically, in c584>TrpA1 flies, increasing temperature from 18°C to 30°C for 12-24 hours decreased sleep (Fig. 2a-b) and subsequently worsened seizure severity (Fig. 2d) predominantly driven by a prolongation of the convulsive phase (as detailed in *Supplementary Methods* flies were brought to room temperature for 20 minute prior to seizure testing) (Ext. Data Fig. 9E-I). After returning the temperature to 18°C, c584>TrpA1 flies exhibited increased nighttime rebound sleep (Fig. 2c) and prolonged sleep bouts (Ext. Data Fig. 9a-d), consistent with a homeostatic sleep recovery. In contrast, Tdc2>TrpA1 is known to lead to sleep loss without homeostatic rebound sleep^29^. In Tdc2>TrpA1 flies, the same temperature shift also reduced sleep (Fig. 2e-f), but here there were minimal effects on seizures (Fig. 3h and Ext. Data Fig. 9n-r). After the temperature was shifted back from 30°C to 18°C, Tdc2>TrpA1 flies continued to demonstrate decreased sleep and no indication of homeostatic sleep rebound or consolidation (Fig. 2g and Ext. Data Fig. 9j-m) as previously described^29^. In conjunction with the effects of sleep need on spontaneous seizures, these thermogenetic experiments indicate that sleepiness, and not sleep amount per se, is the primary driver of seizure severity.

**Fig. 2.**
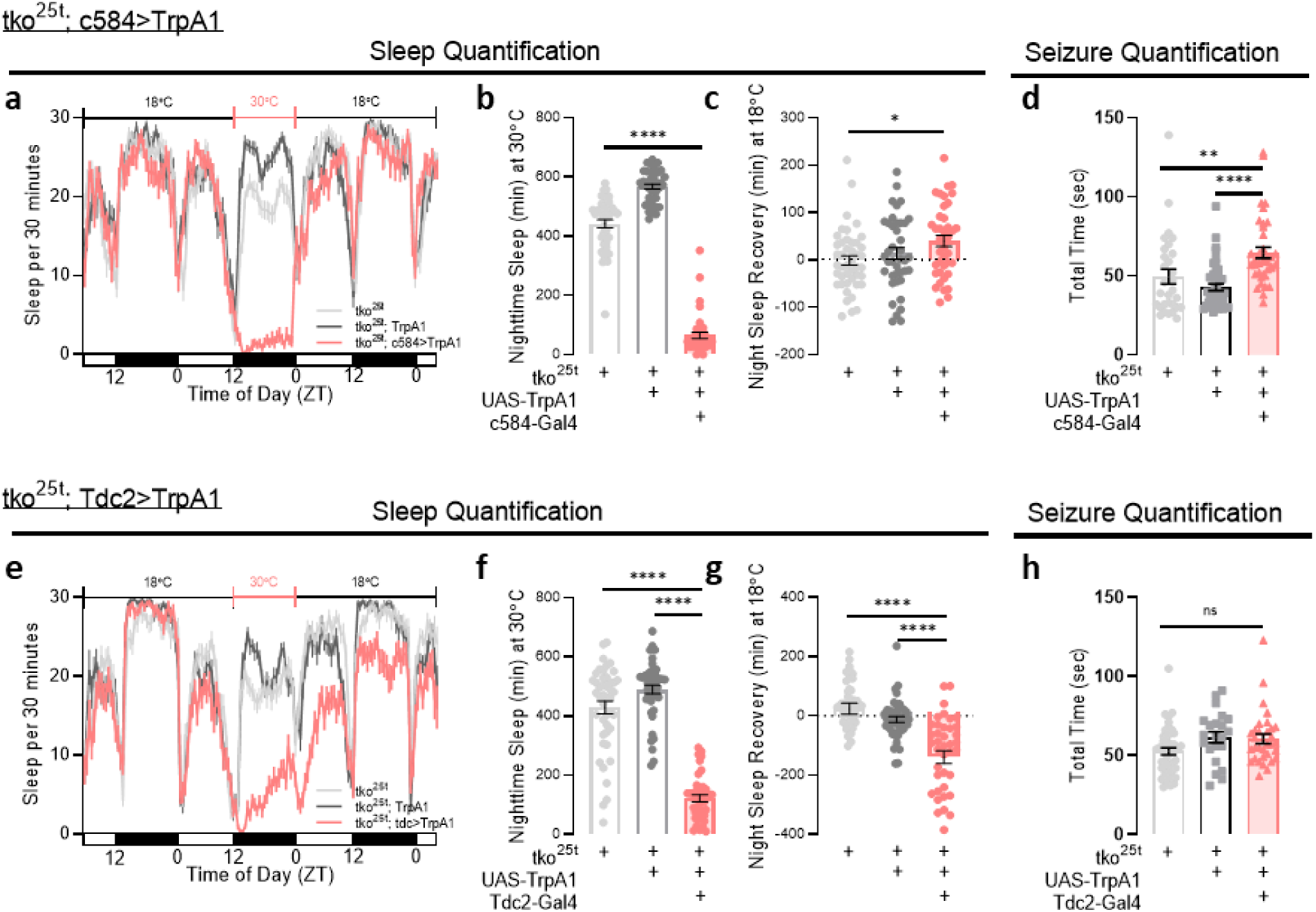
Sleep loss associated with increased sleep need exacerbates seizures. **a-c**, *tko*^25t^; c584-Gal4>UAS-TrpA1 flies were maintained at 18°C, shifted to 30°C overnight for 12 hours, then recovered in 18°C. Thermogenetic activation of c584-Gal4>UAS-TrpA1 at 30°C led to sleep loss that recovered back to baseline at 18°C. After returning to 18°C, night sleep was increased as compared to baseline night sleep, consistent with homeostatic sleep rebound. n=40- 43 flies/condition. **d**, After thermogenetic activation of *tko*^25t^; c584-Gal4>UAS-TrpA1 for 24 hours, total seizure times are prolonged. n=30-45 flies/condition. **e-g**, *tko*^25t^; Tdc2-Gal4>UAS- TrpA1 flies were maintained at 18°C, shifted to 30°C overnight for 12 hours, then recovered in 18°C. Thermogenetic activation of Tdc2-Gal4>UAS-TrpA1 at 30°C led to sleep loss that persisted even after flies were restored to 18°C. There was no homeostatic sleep rebound after temperature was returned to 18°C. n=41-46 flies/condition. **h**, After thermogenetic activation of *tko*^25t^; Tdc2-Gal4>UAS-TrpA1 for 24 hours, there was no significant difference in total seizure times. n=22-45 flies/condition. One-way ANOVA with Tukey’s multiple comparisons test was used. *p<0.05, **p<0.01, ****p<0.0001. Data are presented as mean values ± SEM.

**Fig. 3.**
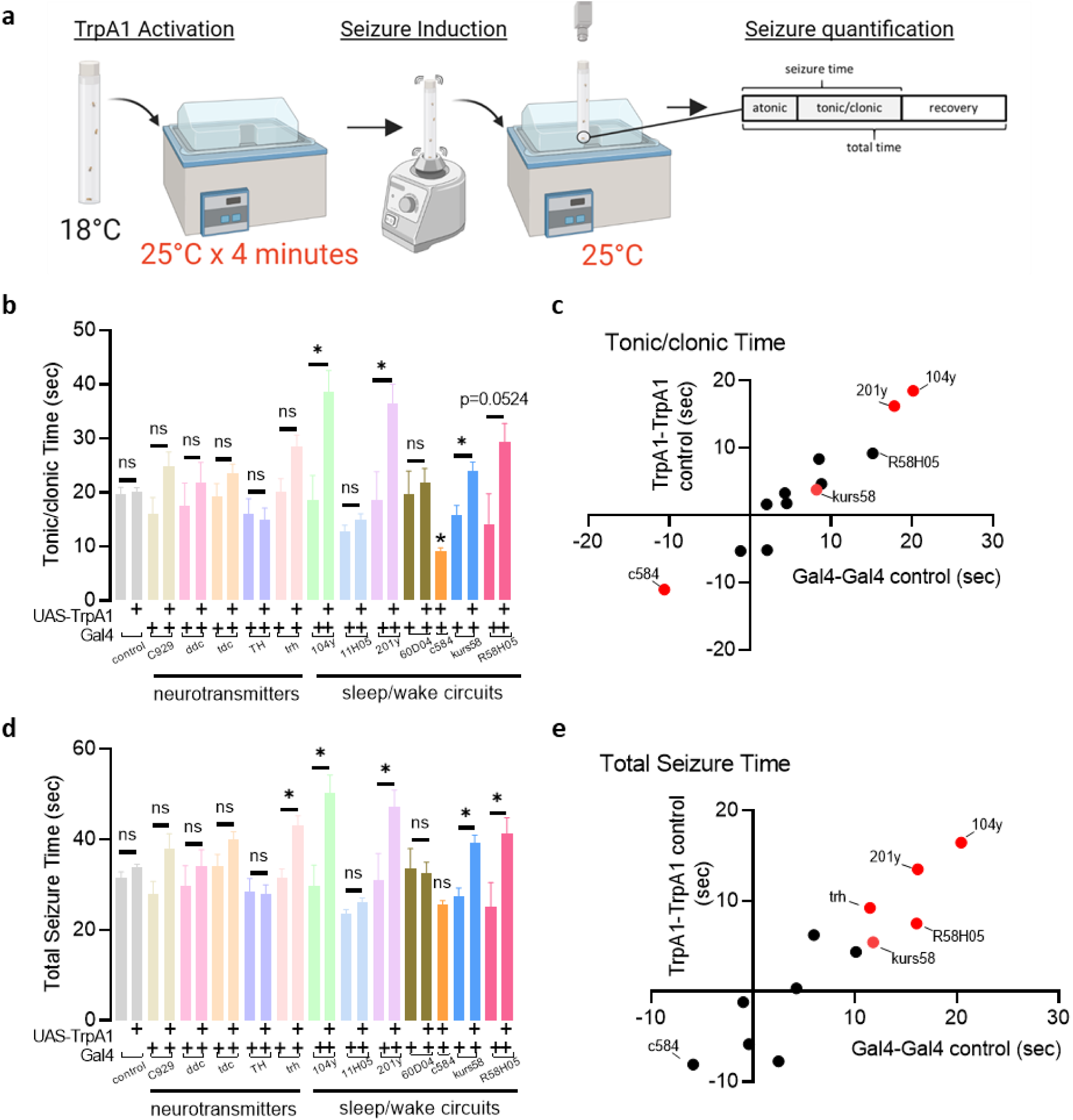
A Gal4 screen reveals activation sleep-promoting circuits worsens seizures. **a**, Experimental protocol depicting flies in the *tko*^25t^ background were raised at 18°C, placed at 25°C for 4 minutes for TrpA1 activation, vortexed for seizure induction, and video recorded for seizures at 25°C. To drive TrpA1, Gal4 drivers were selected for peptidergic (C929-Gal4), serotonergic/dopaminergic (Ddc-Gal4), octopaminergic (Tdc2-Gal4), dopaminergic (TH-Gal4), serotonergic (trh-Gal4), dorsal fan-shaped body (104y-Gal4), wake-promoting (11H05-Gal4, 60D04-Gal4, c584-Gal4), mushroom body (201y-Gal4), pars intercerebralis (kurs58-Gal4), and ellipsoid body (R58H05-Gal4) neurons. **b-c**, Tonic/clonic times were prolonged upon driving of TrpA1 with 104y-Gal4, 201y-Gal4, kurs58-Gal4, and improvement with c584-Gal4. **d-e**, Total seizure times demonstrate worsening of seizures upon driving of TrpA1 with 104y-Gal4, 201y- Gal4, kurs58-Gal4, R58H05-Gal4, and trh-Gal4. n=4-77. Two-group two-tailed t-test. *p<0.05. Data are presented as mean values ± SEM.

### Sleep-promoting circuits modulate seizures

The above data indicate that sleep need might have direct effects on seizure severity independent of sleep amount. To test this, in the *tko*^25t^ background we expressed UAS-TrpA1 in 12 cellular populations or circuits that are known to play a role in wakefulness or sleep. Broad Gal4 drivers were selected for the purposes of screening. Flies were raised at 18°C to minimize TrpA1 activation then brought to 25°C for TrpA1 activation (Fig. 3a). This activation temperature was selected because it adequately increases TrpA1 conductance^30^ and also avoids direct effects of temperature on bang-sensitive seizures in the *tko*^25t^ background^31^. After bang-sensitive seizure induction, flies were then returned to 25°C for video recording and seizure quantification (Fig. 3a). Importantly, in these experiments, discrete cell populations and circuits involved in the sleep/wake cycle were activated, without allowing sufficient time for sleep amount to increase or decrease. This allowed us to test direct effect of the cells/circuits of interest on seizure severity independent of sleep behavior.

Using this paradigm, we found that acute direct activation of sleep-promoting circuits worsened seizure severity. As compared to genetic controls, activation of 104y>TrpA1 (expressed in the dorsal fan-shaped body (dFB)), 201y>TrpA1 (expressed in the mushroom body (MB) as well as 50-100 non-MB cells per hemisphere)^32^, and kurs58>TrpA1 (expressed in the pars intercerebralis (PI)) prolonged the tonic/clonic phase of seizures (Fig. 3b-c). The dorsal fan-shaped body, mushroom body, and pars intercerebralis each have known sleep-promoting functions, as previously reviewed^33^. In addition to drivers expressed in the dFB, MB, and PI, R58H05>TrpA1, expressed in the ellipsoid body (EB), another sleep-promoting region of the brain, increased total seizure times (Fig. 3d-e). While c584>TrpA1 activation reduced tonic/clonic times (note that this protocol differed from the one above in not allowing sleep decrease), other wake-promoting drivers including 11H05>TrpA1 and 60D04>TrpA1 were not protective (Fig. 3b-c). The only neurotransmitter/neuromodulator group that worsened seizure severity upon acute activation were *trh*-expressing serotonergic neurons (Fig. 3d-e). None of the activated populations prolonged recovery times (Ext. Data Fig. 10c), suggesting that other brain regions may be involved in regulation of postictal seizure recovery. In summary, these data demonstrate that acute activation of brain regions that encode sleep need, including the dFB and MB, worsens seizures, even with no change in total sleep amount.

### Sleep loss increases activity of sleep-promoting circuits

These thermogenetic studies implicating the activity of sleep-promoting circuits in seizure severity suggested that sleep-promoting circuits change activity of other brain regions. To address this question, we investigated the effect of sleep loss on brain wide activity by pan-neuronally expressing UAS-CaLexA, a genetically-encoded, fluorescent calcium indicator as a surrogate readout for brain activity. After maintenance of flies on caffeine for 48 hours to restrict sleep, we dissected whole brains and measured endogenous CaLexA signal. Consistent with previous findings, we found increased intracellular calcium in the dFB and EB^7, 9^, as well as the MB (Ext. Data Fig. 11b-d). Incredibly, beyond regions of the brain known to be sleep-promoting, we found that sleep loss induced by caffeine led to brain-wide increases in intracellular calcium (Ext. Data Fig. 11a), indicating that brain-wide activity may be increased in the setting of sleep loss. These effects were not specific to caffeine, as (1) mechanical sleep deprivation and (2) assessment at the end of the day when sleep pressure is high (ZT12), also led to increased calcium in the MB and EB (Ext. Data Fig. 12a-c). These data demonstrate that sleep loss leads to increased activity of sleep-promoting circuits, and this correlates with increased brain-wide activity.

### Circuits encoding sleepiness can toggle seizures

We next asked if direct activation of sleep-promoting circuits increases the likelihood and severity of spontaneous seizures, as it does for bang-sensitive seizures. To achieve broad control of sleep-promoting circuits, we turned to our chronic video tracking paradigm and expressed UAS-csChrimson under control of 23E10-Gal4, which is expressed in the dFB and VNC^7, 8, 15, 16^, as well as 201y-Gal4, expressed in the MB and 50-100 non-MB cells per hemisphere. Flies were raised in the dark, entrained in blue light and fed ATR for 48 hours, then placed into individual wells of a 24-48-well plate containing picrotoxin with stimulation of UAS-csChrimson using red light for the first 5 minutes of every hour (Ext. Data Fig. 13a). Experimental flies and genetic controls were also fed caffeine to test if sleep restriction increases seizures by increasing activity of these sleep-promoting regions, in which case it would have little additional effect, or if it acts synergistically with activation of sleep-promoting regions. As expected, caffeine treatment of genetic controls (23E10-Gal4, 201y-Gal4 + caf) led to sleep loss (Ext. Data Fig. 13b, d) and more frequent and severe seizures (Ext. Data Fig. 13c, f, g).

Prolonged activation of sleep-promoting cells with csChrimson (23E10-Gal4, 201y- Gal4>csChrimson) led to increased sleep (Ext. Data Fig. 13e). However, prolonged activation of sleep-promoting cells in the absence of caffeine did not worsen seizure severity (Ext. Data Fig. 13c, f, g). This is likely because although sleep need was induced, flies had an opportunity to dissipate their sleep need by sleeping more (Ext. Data Fig. 13b, e), thereby reducing their seizure risk. In an effort to induce sleepiness without allowing flies sufficient time to dissipate their sleepiness by sleeping more, we attempted two additional csChrimson stimulation protocols: (1) red light 2 minutes on and 3 minutes off, and (2) red light on continuously. The former protocol enhanced sleep of 23E10-Gal4, 201y-Gal4>csChrimson flies even more, while the constant red-light exposure in the latter protocol disrupted seizure initiation even in control conditions for unclear reasons (data not shown). In contrast, when 23E10-Gal4, 201y-Gal4>csChrimson flies were fed caffeine, there was no similar opportunity to dissipate sleepiness given continued exposure to caffeine (Ext. Data Fig. 13b, d, e); this led to a non-additive increase in seizure frequency and severity, similar to genetic controls fed caffeine (Ext. Data Fig. 13c, f, g). Importantly caffeine treatment of 23E10-Gal4, 201y-Gal4>csChrimson flies led to seizures, including lethal seizures, even though there was increased sleep per fly in the 3 hours preceding seizures as compared to genetic controls (23E10-Gal4, 201y-Gal4 + caf) (Ext. Data Fig. 14c-d). This indicates that aberrant hyperactivation of two brain regions that encode sleep need, marked by 23E10 and 201y drivers, worsens seizures, even if individual flies increase total sleep amount.

If hyperactivation of sleep-promoting circuits increases seizure severity (Ext. Data Fig. 6c, f), then this raised the exciting possibility that inactivation of sleep-promoting circuits might be a strategy to decrease seizure risk. For inhibition, we expressed UAS-GtACR1, a light-sensitive anion channelrhodopsin in sleep-promoting cells using 23E10-Gal4 and 201y-Gal4. Similar to the protocol for csChrimson activation, we raised flies in the dark then entrained flies for 48 hours in red light with ATR, then placed them into 24-48-well plates containing picrotoxin (Fig. 4a). While caffeine treatment of genetic controls (23E10-Gal4, 201y-Gal4 + caf) led to decreased sleep (Fig. 4b, d, e) and increased seizures (Fig. 4c, f, g), experimental 23E10-Gal4, 201y- Gal4>GtACR1 flies fed caffeine showed decreased sleep (Fig. 4b, d, e) and also decreased seizures (Fig. 4c, f, g). Inhibition of sleep-promoting cells in 23E10-Gal4, 201y-Gal4>GtACR1 flies in vehicle conditions also led to decreased sleep, comparable to caffeine treatment, and decreased seizures (Fig. 4b, d, e). Reminiscent of the effects of Tdc2>TrpA1 noted above, this reduced sleep amount is presumably associated with decreased sleep need secondary to direct inactivation of sleep-promoting cells with GtACR1. These optogenetic experiments provide direct evidence that the effects of sleep need on seizure risk can be dissociated from the effects of sleep amount.

**Fig. 4.**
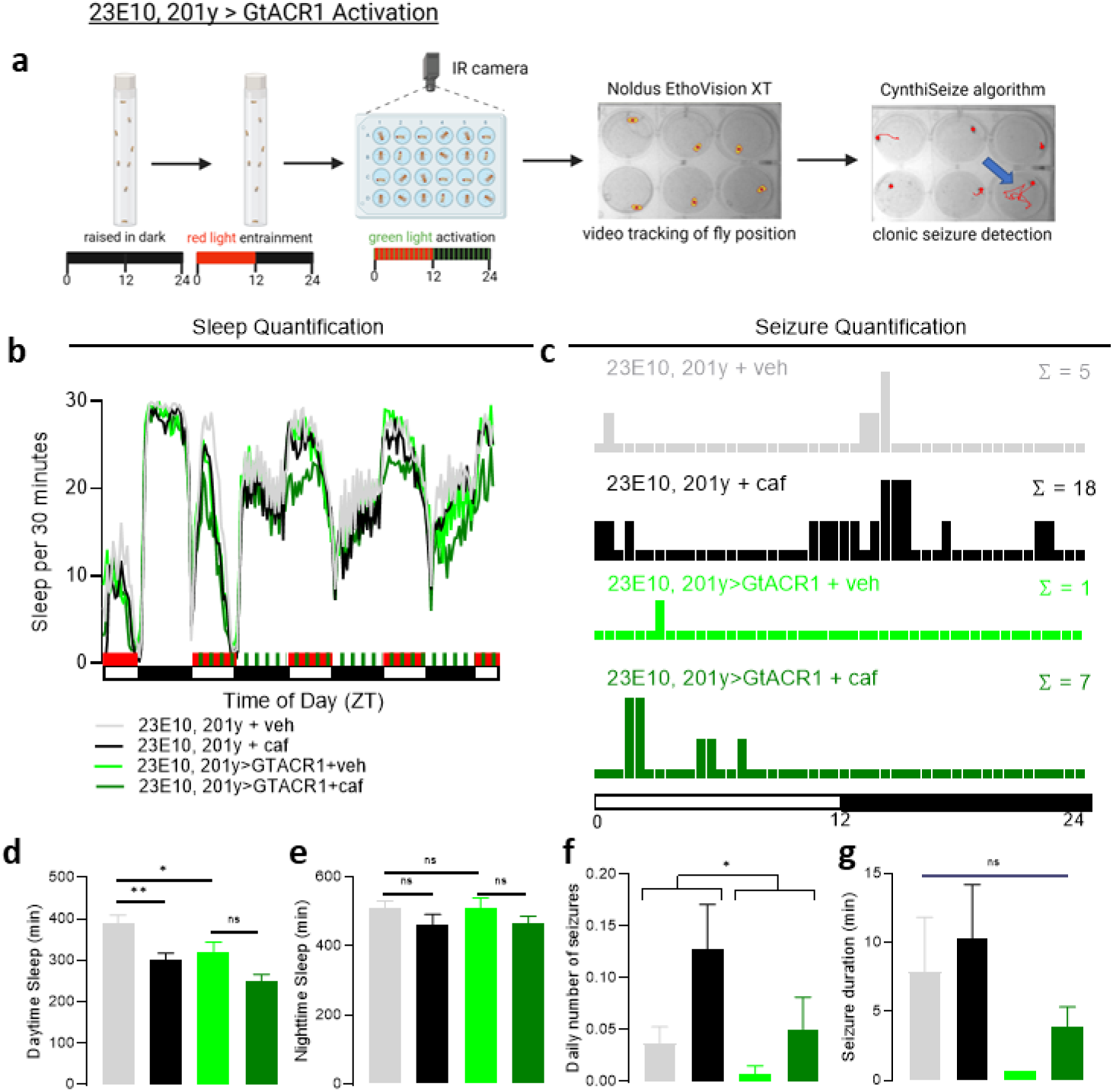
Inhibition of sleep-promoting circuits protects against seizures in the setting of sleep loss. **a**, Experimental protocol showing flies were raised in darkness, entrained in red light, then placed into 24- or 48-well plates for chronic video monitoring with green light stimulation. Fly positions over time were converted into XY coordinates, and a “CynthiSeize” algorithm was developed to identify seizures. **b**, **d**, **e**, Sleep quantification after 23E10-Gal4, 201y- Gal4>GtACR1 activation. Caffeine and 23E10-Gal4, 201y-Gal4>GtACR1 decrease sleep. n=34 flies/condition. **c**, **f**, **g**, Seizure quantification after 23E10-Gal4, 201y-Gal4>GtACR1 activation. Caffeine increases seizure frequency and this is blocked with 23E10-Gal4, 201y-Gal4>GtACR1 activation. n=34 flies/condition. One-way ANOVA with Dunnett’s T3 multiple comparisons adjustment, Kruskal-Wallis test with Dunn’s multiple comparisons adjustment, negative binomial model with Wald test (for daily number of seizures), or mixed effects model (for seizure durations) was used. *p<0.05, **p<0.01, ***p<0.001. Data are presented as mean values ± SEM.

### Loss of 5HT1A mediates increased seizures

To understand how sleep-promoting circuits might become hyperactive to increase seizure burden, we performed RNA sequencing of the dFB with and without sleep deprivation. Given the ability of the dFB to affect whole-brain activity, we focused our attention on transcripts encoding neurotransmitter or neuromodulator receptors and rate limiting enzymes (Fig. 5a). Of these transcripts, only one significantly changed after sleep restriction: 5HT1A. The serotonin 5HT1A receptor is a G protein-coupled receptor that couples with inhibitory Gi proteins as previously reviewed^34^. Downregulation of 5HT1A in the dFB after sleep restriction (Fig. 5a) would be predicted to decrease inhibitory signaling, thereby leading to increased activity of the dFB and enhanced sleep pressure.

**Fig. 5.**
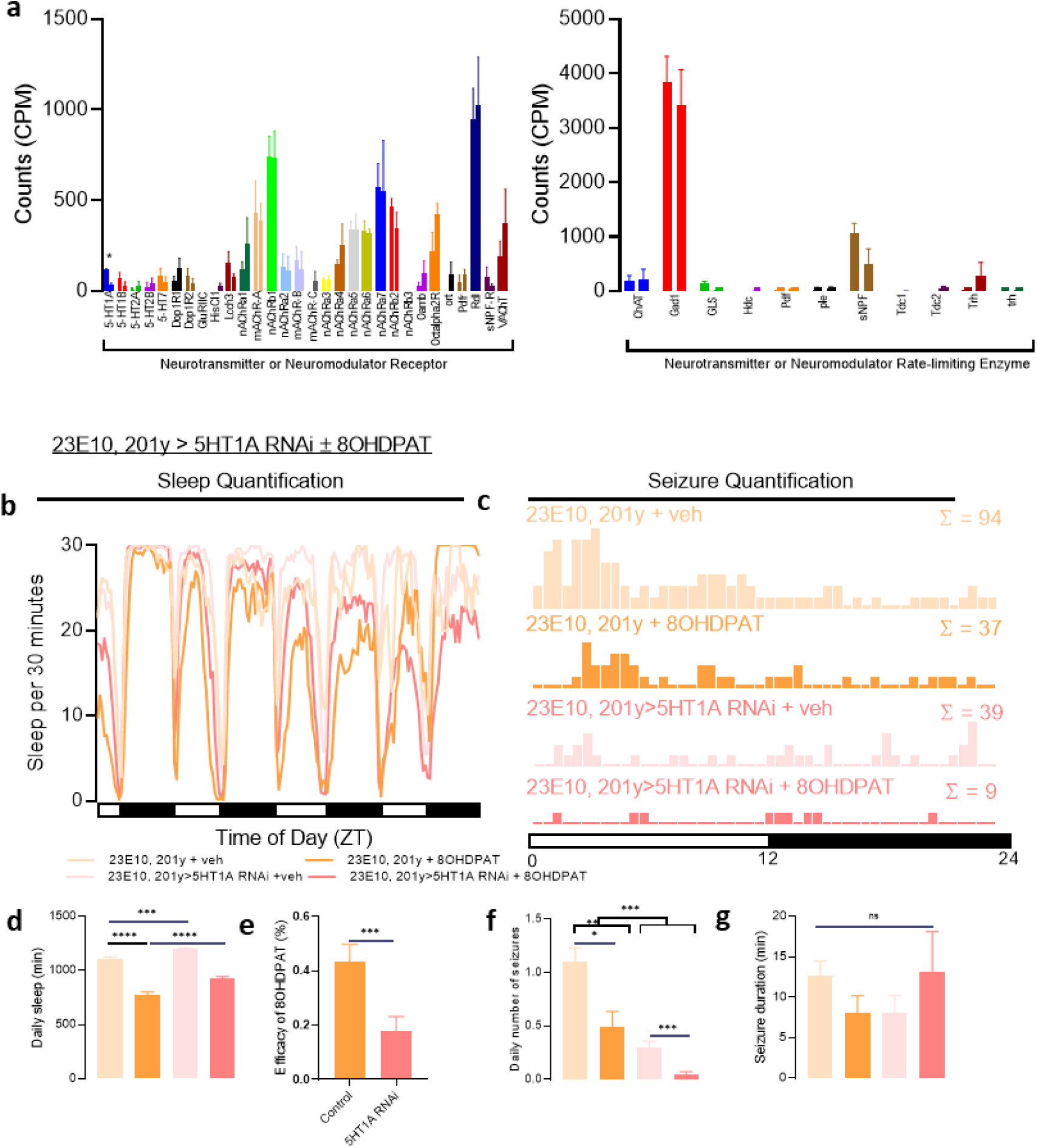
5HT1A is downregulated after sleep loss, and this is sleep-promoting. **a**, Transcriptomic analysis of the dorsal fan-shaped body after sleep-restriction reveals downregulation of 5HT1A, but not other neurotransmitter receptors or rate-limiting enzymes. **b**, **d**, Sleep quantification after addition of 8-OH-DPAT, a selective 5HT1A receptor agonist, reveals decreased sleep, consistent with the inhibitory effects of 5HT1A on the activity of sleep-promoting centers. Conversely, 23E10-Gal4, 201y-Gal4> 5HT1A RNAi-mediated downregulation of 5HT1A in sleep-promoting centers leads to increased sleep. n=46 flies/condition. **e**, The effect of a 5HT1A agonist on daily sleep loss is decreased after downregulation of 5HT1A in sleep-promoting centers. **c**, **f**, **g**, Seizure severity is reduced after both 5HT1A agonism and 5HT1A downregulation. One-way ANOVA with Dunnett’s T3 multiple comparisons test, paired two-tailed t-test, negative binomial model with Wald test (for daily number of seizures), or mixed effects model (for seizure durations) was used. *p<0.05, ***p<0.001, ****p<0.0001. Data are presented as mean values ± SEM.

We hypothesized that activation of 5HT1A in sleep-promoting circuits would decrease sleep. To test this hypothesis, we used our chronic video tracking platform (Fig. 1a) and induced seizures with picrotoxin. As hypothesized, genetic control flies (23E10-Gal4, 201y-Gal4) fed 8-OH-DPAT, a selective 5HT1A receptor agonist, exhibited decreased sleep (Fig. 5b, d). Although there was decreased sleep, there was a decrease in seizure frequency (Fig. 5c, f), presumably because 5HT1A-mediated signaling increases inhibition of sleep-promoting cells, thereby decreasing both sleep and sleep need. To better understand the site of action of 8-OH-DPAT, we knocked down expression of 5HT1A in sleep-promoting cells using RNAi (23E10-Gal4, 201y- Gal4>5HT1A RNAi). Knockdown of 5HT1A in sleep-promoting cells caused an increase in sleep (Fig. 5b, d). This is consistent with the RNAseq results (Fig. 5a), as sleep loss leads to loss of 5HT1A expression, thereby driving increased activity of the dFB and leading to more sleep. The increased sleep resulting from loss of 5HT1A expression in the dFB and MB would lead to dissipation of sleep need. Consistent with this possibility, we found loss of 5HT1A expression led to decreased seizure risk (Fig. 5c, f, g, Ext. Data Fig. 16b). When 8-OH-DPAT was fed to 23E10-Gal4, 201y-Gal4>5HT1A RNAi flies, it was significantly less effective at reducing sleep, indicating the effects of 5HT1A agonism is mostly occurring through its expression in sleep-promoting cells (Fig. 5d, e). In summary, these results demonstrate that the loss of 5HT1A- mediated inhibitory signaling promotes sleep. We also find that agonism of 5HT1A can be used to decrease sleep pressure and protect against seizures.

Given that increased 5HT1A signaling reduced sleep pressure and protected against seizures, we asked if a clinically available, Food and Drug Administration (FDA)-approved medication might be able to reverse the effects of sleepiness on seizures. Buspirone has high affinity for the serotonin 5HT1A receptor, whereby it acts as a partial agonist, and is clinically approved to treat mood disorders. Consistent with the inhibitory effects of 5HT1A on sleep-promoting circuits described above, a side-effect of buspirone is insomnia^35^. We fed wild-type flies buspirone and caffeine to determine if buspirone could decrease sleep need through 5HT1A agonism, thereby reducing the increased seizure burden associated with sleep loss. Buspirone did not have an effect on baseline sleep amount as compared to 8-OH-DPAT (Ext. Data Fig. 15a, c, d). As previously seen, caffeine led to decreased sleep and increased seizures (Ext. Data Fig. 15a-f). The addition of buspirone to caffeine did not significantly affect caffeine-induced sleep loss (Ext. Data Fig. 15a, c, d), but it did decrease seizure risk (Ext. Data Fig. 15b, e). Thus, we find that buspirone, an FDA-approved 5HT1A agonist, is able to reverse the increased seizure burden occurring after sleep restriction.

## Discussion

Our data suggest that after sleep loss, the primary driver of worsened seizures is sleep need, and not sleep amount. We find that manipulations that sustain activity of sleep-promoting centers increase seizure frequency. Importantly, the converse is also true: if excitation of sleep-promoting centers is decreased, then seizures are less severe, even in the setting of sleep loss. We propose that targeting neural correlates of sleepiness can improve seizure control (Ext. Data Fig. 17).

In considering the effects of sleep on seizures, there are multiple variables to account for, including, but not limited to, (1) sleep history, which is the total sleep amount prior to an event of interest, (2) sleepiness, which is the drive to sleep more, (3) sleep or wake status at the time of an event of interest, and (4) how recently a state change (wake → sleep or sleep → wake) occurred.

Our video tracking platform allowed for quantification of these variables while simultaneously monitoring for seizures in *Drosophila.* We present multiple lines of evidence demonstrating that the drive to sleep has a greater impact on seizure burden than sleep amount. These results provide an explanation to a discrepancy in clinical neurology: while individuals with epilepsy identify decreased sleep as a trigger for seizures^2, 3^, objective measurement of sleep amount in individuals with epilepsy demonstrates a poor correlation with seizure risk^36^. Given sleep need is variable between individuals^37^, our data suggest that increasing sleep quantity is not key to preventing seizures, but rather providing sufficient rest to minimize sleep drive decreases seizure risk.

If sleep loss leading to sleepiness worsens seizures, then this raises the interesting possibility that enhancing sleep to dissipate sleepiness might be protective against seizures. Therefore, we attempted two manipulations to increase total sleep: gaboxadol feeding (Ext. Data Fig. 1) and *nemuri* overexpression (Ext. Data Fig. 4). While both manipulations increased sleep, there were seemingly opposite effects on seizures; gaboxadol worsened seizures (Ext. Data Fig. 1i) while *nemuri* overexpression was strongly protective (Ext. Data Fig. 4d-i). We propose this can be explained by variable effects on sleepiness. The sleep induced by gaboxadol is known to be different from spontaneous sleep as measured by local field potentials in *Drosophila*^38^ and murine electroencephalography^39^. We suggest that the sleep induced by gaboxadol is not the same as spontaneous restorative sleep and is not able to fully dissipate sleep pressure. Therefore, ongoing sleep pressure led to the worsened seizures we observed after gaboxadol treatment. In contrast, *nemuri* overexpression is known to increase both sleep amount and promote deeper sleep depth^22^. In this deep sleep state induced by *nemuri* overexpression, sleep need may not accumulate suggesting a protective effect against seizures.

Given that seizures are not generated in sleep-promoting circuits, we sought to understand how sleep need affects activity across the brain. Using intracellular calcium concentration as a proxy for neuronal activity, we confirmed that elevated sleep pressure is associated with increased activity of sleep-promoting regions of the brain (Ext. Data Fig. 11, Ext. Data Fig. 12), consistent with previous findings^7–9^. We find that this increased activity is not isolated to sleep-promoting circuits and occurs more broadly throughout the brain (Ext. Data Fig. 11a). Flies awake for 5-8 hours exhibit increased intraneuronal calcium concentrations with increased reactivity to external stimuli, possibly driven by an increase size and number of synapses^6, 40, 41^. Similar widespread increases in cortical excitability are known to occur in rodent models^42^ and people^43–46^. We propose that these broad increases in neuronal excitability across the brain drive elevated seizure susceptibility. This is supported by mammalian models demonstrating that sleep restriction increases seizure risk and severity^47–49^. Taken together, our data are consistent with a model wherein activity of the whole brain increases with accumulating sleep pressure.

The use of serotonergic drugs to treat epilepsy has received growing attention since the FDA approved fenfluramine in 2020 to treat seizures associated with Dravet Syndrome. Fenfluramine is thought to act through 5-HT1D- and 5-HT2C-agonism^50^ to improve seizure control. We find that use of 5HT1A agonists can be used to suppress seizures that are worsened with sleepiness in *Drosophila* (Ext. Data Fig. 15). We identify an FDA-approved agonist, buspirone, to decrease seizures sensitive to sleepiness. Broad activation of serotonergic neurons with trh-Gal4>TrpA1 led to worsened seizures (Fig. 3d-e), demonstrating that broadly promoting serotonergic tone may not be helpful and further emphasizing the importance of 5HT receptor specificity. Despite the growing number of anti-seizure medications clinically available, about one-third of individuals with epilepsy continue to have seizures refractory to medical therapy. Targeting non-conventional mechanisms, including the effects of sleepiness on seizures, may offer improved clinical management.

## Methods

Adult male mated flies were used, 5-10 days post-eclosion. Flies were raised in *Drosophila* incubators with a 12-hour light:12-hour dark light cycle at 25° and approximately 60-65% relative humidity. Flies were maintained on a standard cornmeal-molasses diet, including 64.7g/L cornmeal, 27.1g/L dry yeast, 8g/L agar, 61.6mL/L molasses, 10.2mL/L 20% tegosept, and 2.5mL/L propionic acid. A detailed description of all experimental methodology is located in the Supplementary information.

## Data availability

Our transcriptomics dataset of control versus sleep-deprived neurons from the dorsal fan-shaped body will be deposited online. Other data and additional information required to reanalyze data reported in this paper is available from the lead contact upon request.

## Code availability

Our scripts to detect sleep and seizures from video tracking data will be deposited to the repository on GitHub (https://github.com/cthsu86).

## Acknowledgments

We thank Kiet Luu for assistance with fly maintenance. We thank members of the laboratory for helpful discussions. Schematics were created with Biorender (biorender.com). The article is subject to HHMÌs Open Access to Publications policy. HHMI lab heads have previously granted a nonexclusive CC BY 4.0 license to the public and a sublicensable license to HHMI in their research articles. Pursuant to those licenses, the author-accepted manuscript of this article can be made freely available under a CC BY 4.0 license immediately upon publication. The content is the sole responsibility of the authors and does not necessarily represent the official views of the National Institutes of Health.

## Author contributions

Conceptualization: VAC, AS; Methodology: VAC, CH, AS; Investigation: VAC, CH, YL, CS, HS, SK, JS, ZY, GG, MEP, AS; Visualization: VAC; Funding acquisition: VAC, AS; Supervision: VAC, AS; Writing – original draft: VAC, AS; Writing – review & editing: VAC, CH, YL, GG, MEP, AS

## Additional information

Supplementary methods, discussion, references, and movies are available for this manuscript. Correspondence and requests for data or materials should be addressed to Amita Sehgal, PhD at

## Extended data

**Ext. Data Fig. 1.**
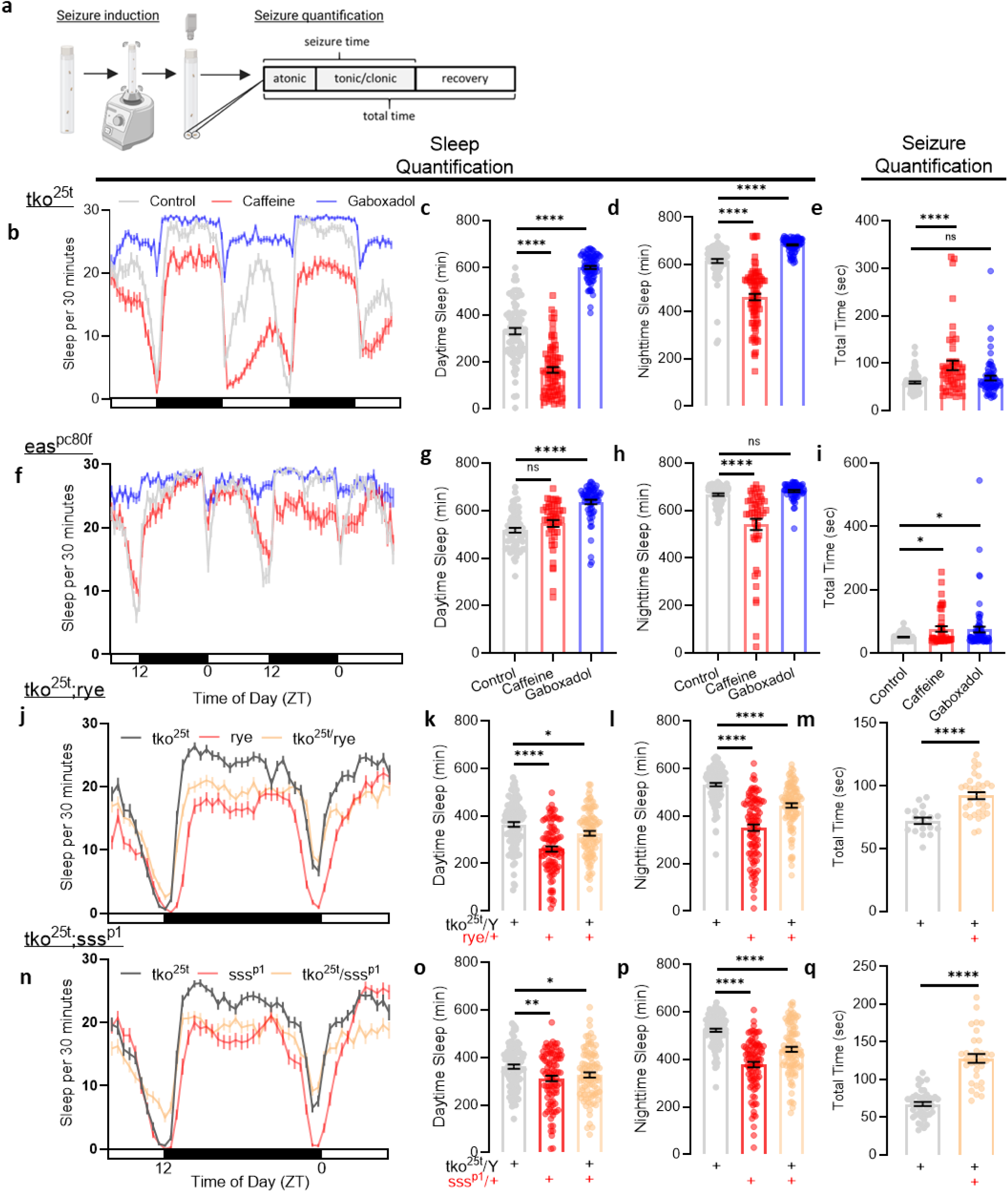
Sleep loss leads to more severe induced seizures in bang-sensitive *tko*^25t^ and *eas^pc^*^80f^ mutant flies. **a**, Experimental protocol depicting seizure induction on vortexer followed by video recording of fly seizures. Seizures are quantified for atonic (“paralysis”), tonic/clonic (“convulsive”), and recovery (“postictal”) phases. “Seizure time” is atonic + tonic/clonic phases. “Total time” is “seizure time” + recovery phase. **b-d**, *tko*^25t^ flies exhibit decreased mean sleep times with caffeine and increased mean sleep times with gaboxadol. n=78 flies/condition. **e**, *tko*^25t^ flies demonstrate prolonged mean seizure duration with caffeine treatment. n=51-74 flies/condition. **f-h**, *eas^pc^*^80f^ flies exhibit decreased mean nighttime sleep times with caffeine and increased mean sleep times with gaboxadol. n=47-62 flies/condition. **i**, *eas^pc^*^80f^ flies demonstrate prolonged mean seizure times with caffeine and gaboxadol. n=43-88 flies/condition. **j-l**, *tko*^25t^*;rye* flies exhibit decreased mean daytime and nighttime sleep times. n=95-96 flies/condition. **m**, *tko*^25t^*;rye* flies demonstrate prolonged mean seizure times. n=18-33 flies/condition. **n-p**, *tko*^25t^*;sss^p^*^1^ flies exhibit decreased mean daytime and nighttime sleep times. n=93-96 flies/condition. **q**, *tko*^25t^*;sss^p^*^1^ flies demonstrate prolonged mean seizure times. n=33-44 flies/condition. Two-group two-tailed t-test or one-way ANOVA with Dunnett’s or Tukey’s multiple comparisons test was used. *p<0.05, **p<0.01, ****p<0.0001. Data are presented as mean values ± SEM.

**Ext. Data Fig. 2.**
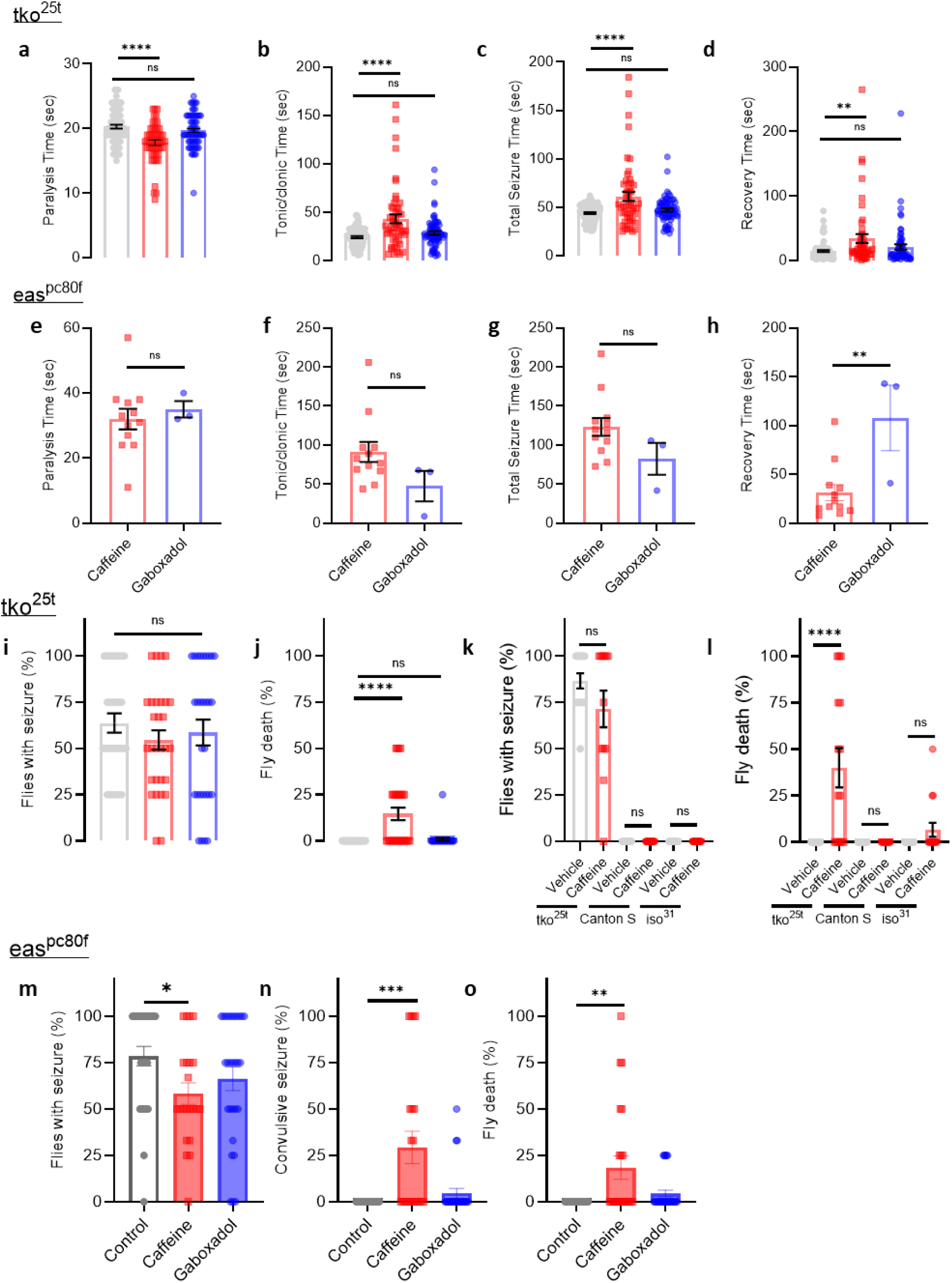
Caffeine leads to more severe induced seizures and death in bang-sensitive mutant flies. **a-d**, Compared to control, *tko*^25t^ flies exhibit decreased mean paralysis time and prolonged tonic/clonic, total seizure, and recovery time with caffeine treatment (*red*). Compared to control, gaboxadol treatment does not significantly change mean seizure times. n = 51-74 flies/condition. **e-h**, In *eas^pc^*^80f^ flies, there is no significant evidence of differences in mean paralysis time, tonic/clonic time, and between caffeine and gaboxadol treatment. Recovery times are significantly prolonged with gaboxadol treatment as compared to caffeine treatment. Vehicle- fed *eas^pc^*^80f^ flies did not exhibit tonic/clonic seizures, therefore no control conditions are listed. n = 43-88 flies/condition. Note there are far fewer tonic/clonic seizures among *eas^pc^*^80f^ flies, and thus the statistical power to detect differences is lower than for a-d. **i-j**, We did not observe differences in induced seizure likelihood after caffeine treatment for 48 hours among *tko*^25t^ flies, but flies were more likely to die. n = 29 vials/condition. **k-l**, *tko*^25t^ flies had comparable numbers of seizures but were more likely to die after caffeine treatment for 60 hours. Canton S and iso^31^ wildtype flies never exhibited seizures and are less likely to die after caffeine treatment for 60 hours. n = 12-15 vials/condition. **m-o**, *eas^pc^*^80f^ flies are less likely to exhibit induced seizures and more likely to die after caffeine treatment for 48 hours. When seizures do occur, *eas^pc^*^80f^ flies are more likely to exhibit convulsive seizures after caffeine treatment. n = 21-28 vials/condition. Two-group two-tailed t-test, unpaired two-tailed t-test with Welch’s correction, one-way ANOVA with Dunnett’s multiple comparisons test, or Kruskal-Wallis with Dunn’s multiple comparisons test (for seizure percentage or fly death percentage) was used. *p<0.05, **p<0.01, ***p<0.001, ****p<0.0001. Data are presented as mean values ± SEM.

**Ext. Data Fig. 3.**
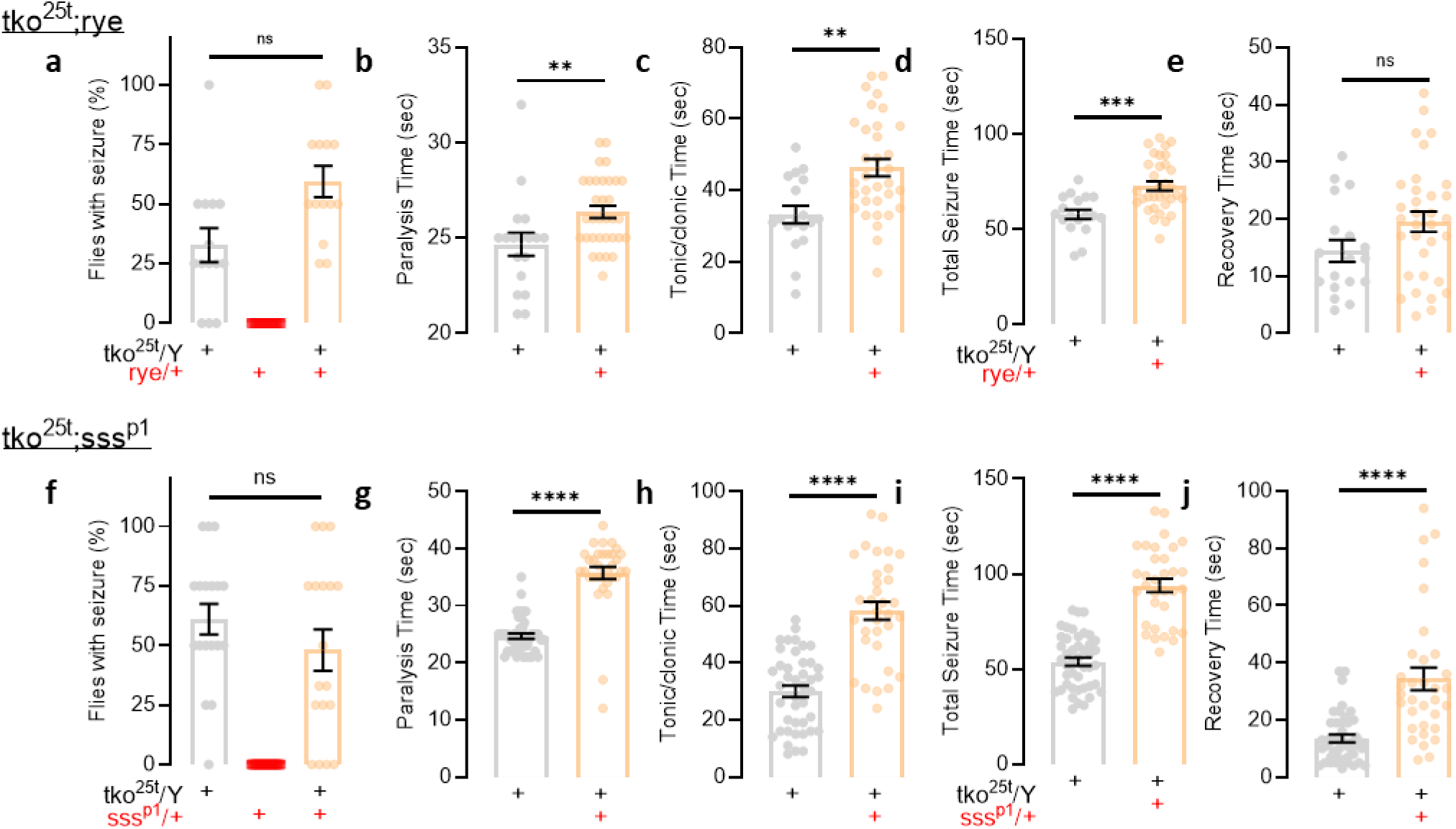
Crossing short-sleeping flies with *tko*^25t^ mutant flies leads to more severe induced seizures. **a-e**, *tko*^25t^*;rye* flies have more severe seizures as compared to mutant *tko*^25t^ flies. n = 14 vials/condition and n = 18-33 flies/condition. **f-j**, *tko*^25t^*;sss^p^*^1^ flies exhibit prolonged seizures as compared to mutant *tko*^25t^ flies. n = 18 vials/condition. n = 33-44 flies/condition. Two-group two-tailed t-test, one-way ANOVA with Dunnett’s or Tukey’s multiple comparisons test, or Kruskal-Wallis with Dunn’s multiple comparisons (for seizure percentage) test was used. **p<0.01, ***p<0.001, ****p<0.0001. Data are presented as mean values ± SEM.

**Ext. Data Fig. 4.**
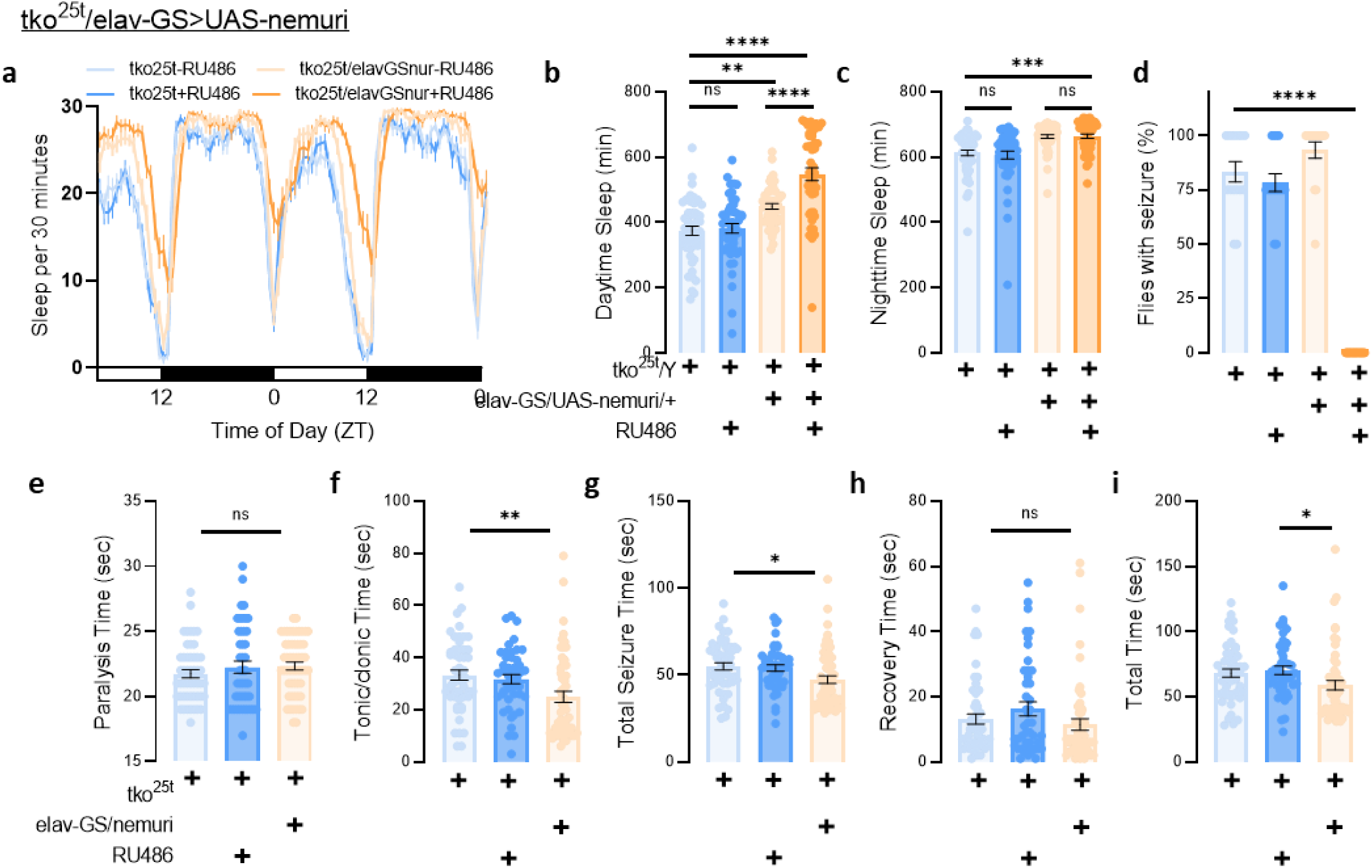
Sleep induction through *nemuri* overexpression leads to decreased seizure likelihood and severity. **a-c**, In *tko*^25t^ mutant flies, overexpression of UAS-*nemuri* using an inducible, pan-neuronal Gal4 driver (*elav*-GeneSwitch, *elav*-GS) by feeding flies the GeneSwitch activator RU486 leads to increased daytime and nighttime sleep n = 45-48 flies/condition. **d-i**, *nemuri* overexpression leads complete suppression of seizures. Even in the absence of the GeneSwitch activator RU486, there are decreased seizure times, likely due to leaky nemuri expression. n = 15 vials/condition. n = 45-55 flies/condition. One-way ANOVA with Dunnett’s multiple comparisons test or Kruskal-Wallis with Dunn’s multiple comparisons test (for seizure percentage) was used. *p<0.05, **p<0.01, ***p<0.001, ****p<0.0001. Data are presented as mean values ± SEM.

**Ext. Data Fig. 5.**
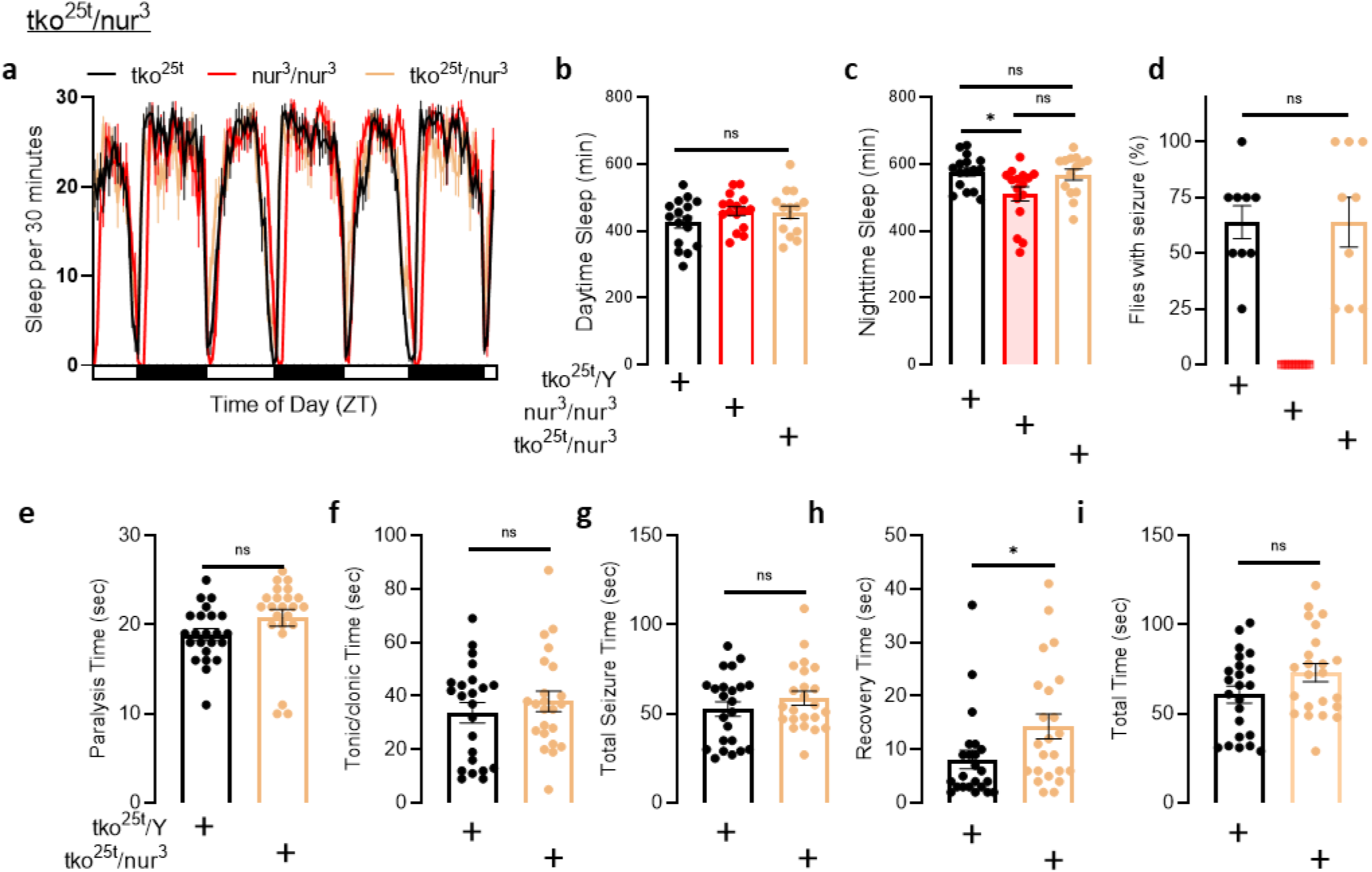
Crossing *tko*^25t^ mutant flies with *nemuri* mutant flies has little effects on sleep or seizure severity. **a-c**, Crossing *tko*^25t^ mutant flies with *nemuri* mutant flies (*nur*^3^) was not associated with significant changes in sleep duration. n = 14-16 flies. **d-i**, There was no significant evidence of a change in seizure likelihood and duration among *tko*^25t^*;nur*^3^ mutant flies. n = 9 vials/condition. n = 23 flies/condition. Two-group two-tailed t-test, one-way ANOVA with Dunnett’s multiple comparisons test, or Kruskal-Wallis with Dunn’s multiple comparisons test (for seizure percentage) was used. *p<0.05. Data are presented as mean values ± SEM.

**Ext. Data Fig. 6.**
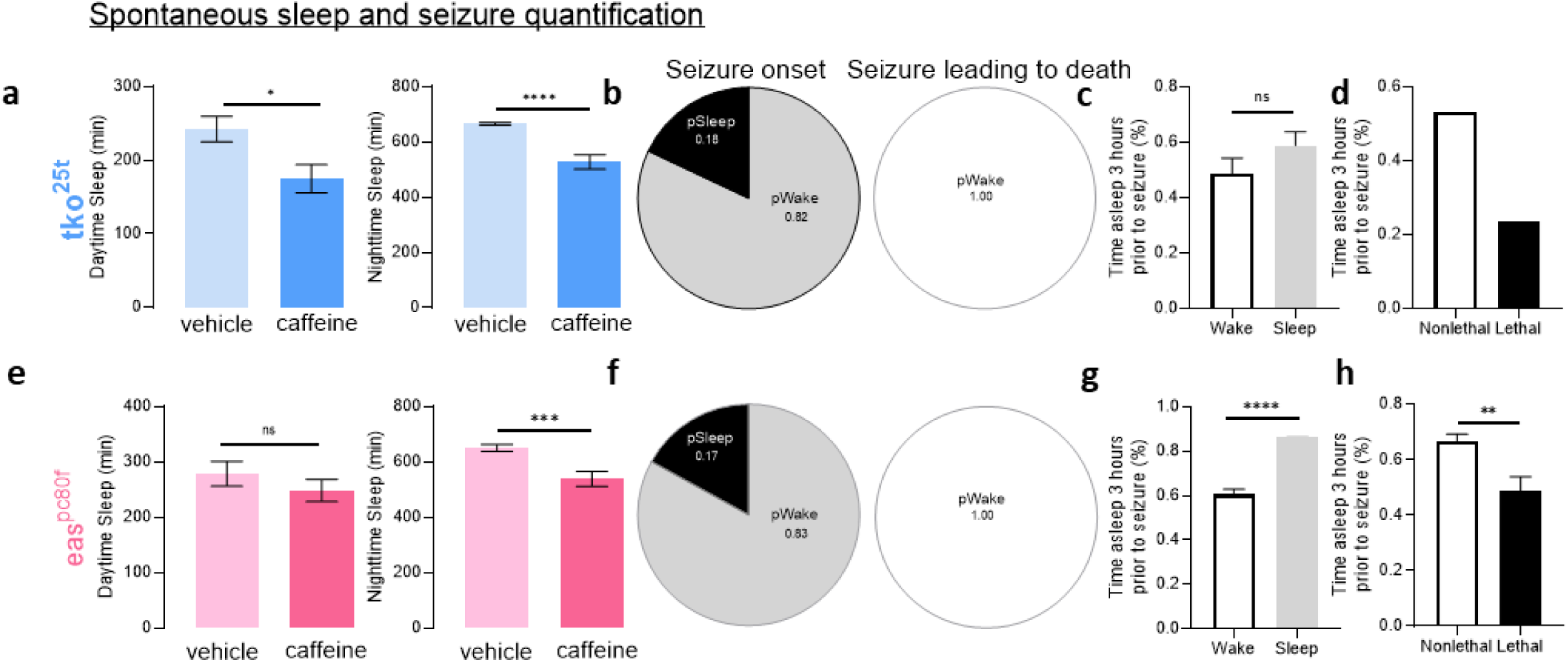
A new protocol for detection of spontaneous seizures in *Drosophila* reveals spontaneous seizures after sleep restriction. **a**, *tko*^25t^ mutant flies treated with caffeine exhibit decreased mean levels of daytime and nighttime sleep as assessed with video tracking. (Lighter colors are with vehicle and darker colors are with caffeine treatment). n = 29-32 flies/condition. **b**, Probability of the fly being in a sleep or wake state at the time of seizure onset for all seizures (left) and seizures that end in death (right). *tko*^25t^ mutant flies more frequently exhibit seizures during wakefulness. Lethal seizures always occur during wakefulness. **c**, Sleep history is similar between *tko*^25t^ mutant flies that have seizures during wakefulness versus those that seize during sleep. n = 2-9 seizures/condition. **d**, Lethal seizures correlate with less time asleep preceding seizures. Error bars not depicted given only a single seizure in *tko*^25t^ mutant flies led to death. n = 1-10 seizures/condition. **e**, *eas^pc^*^80f^ mutant flies treated with caffeine exhibit decreased nighttime sleep as assessed with video tracking. n = 29-32 flies/condition. **f**, *eas^pc^*^80f^ mutant flies more frequently exhibit seizures during wakefulness. Lethal seizures always occur during wakefulness. **g**, Seizures occurring in sleep correlate with a greater percentage of the preceding 3 hours spent in sleep. n = 14-69 seizures/condition. **h**, Lethal seizures correlate with less time asleep preceding seizures in *eas^pc^*^80f^ flies. n = 9-74 seizures/condition. Two-group two-tailed t-test was used. *p<0.05, **p<0.01, ***p<0.001, ****p<0.0001. Data are presented as mean values ± SEM.

**Ext. Data Fig. 7.**
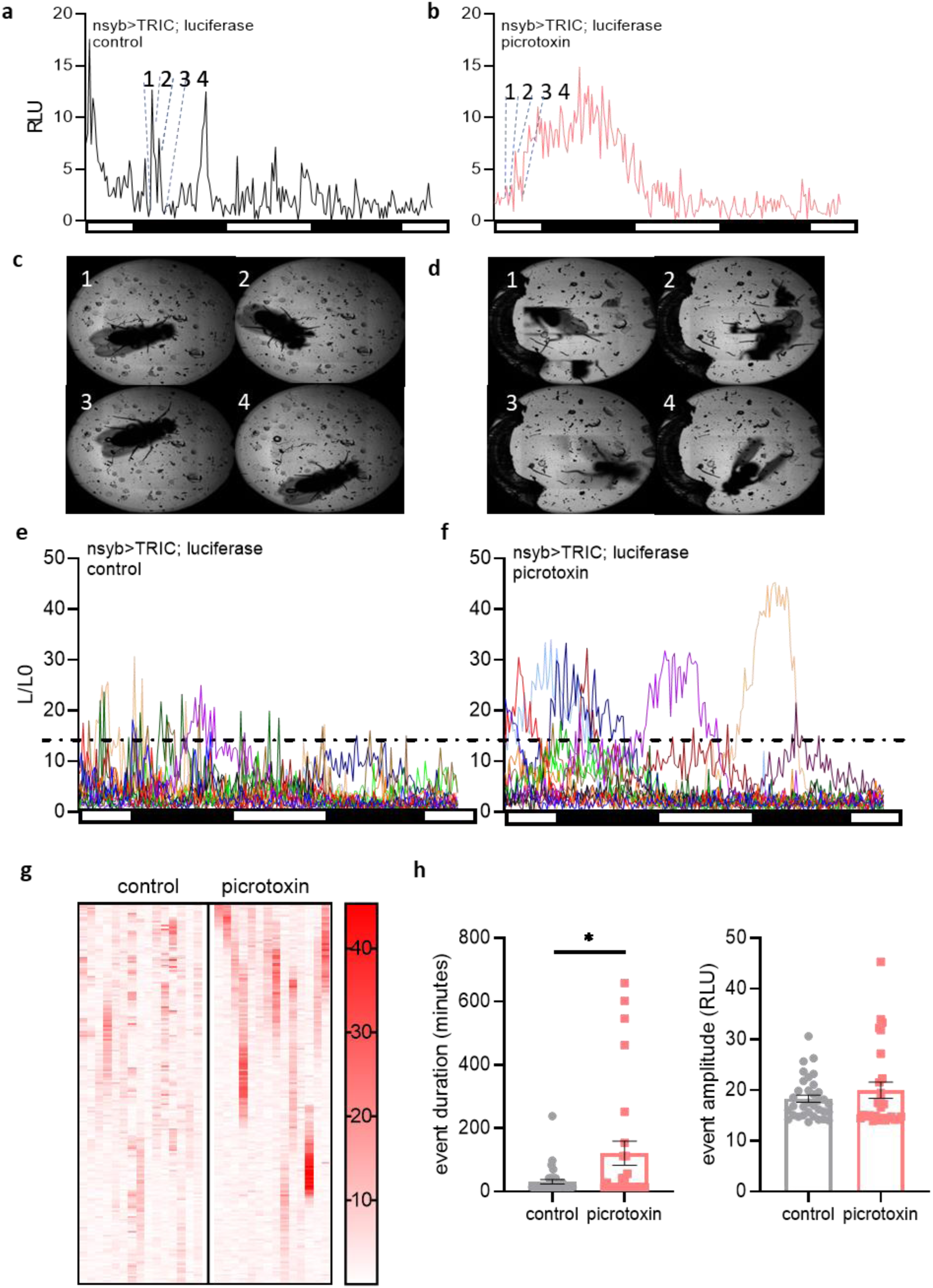
Spontaneous tonic-clonic seizures in *Drosophila* are associated with prolonged, hypersynchronous neuronal activity. nsyb-Gal4>UAS-TRIC-luciferase flies were treated with vehicle or picrotoxin and monitored for 48 hours with imaging of fly posture and detection of luminescence. **a**, **c**, Representative control nsyb-Gal4>UAS-TRIC-luciferase fly treated with vehicle demonstrates that transient spikes in neuronal activity does not correlate with hyperkinetic movements. Timepoints *1, 2, 3,* and *4* in **a** demonstrate a sample spike in luminescence does not correlate with tonic-clonic movements in fly at the same timepoints in **c**. **b, d**, Representative nsyb-Gal4>UAS-TRIC-luciferase fly treated with picrotoxin demonstrates prolonged increases in neuronal activity correlates with hyperkinetic movements. Timepoints *1, 2, 3,* and *4* in **b** demonstrate a prolonged increase in luminescence that correlates with tonic-clonic movements at the same timepoints in **d**. Note that each well containing a fly is constructed from a composite of multiple images of the well. When flies have quick convulsive movements, wings and legs appear blurred, and the fly body appears fragmented. **e-f**, Each colored line depicts an individual fly monitored over two days from a representative experiment. Picrotoxin-treated nsyb-Gal4>UAS-TRIC-luciferase flies exhibit prolonged increases in neuronal activity as measured by luminescence recordings. Recordings are normalized to baseline and presented as L/L0. **g**, Heatmap of same flies as in E-F, but each column demonstrates luminescence over time. Flies treated with picrotoxin exhibit prolonged hypersynchronous neuronal activity. **h**, Quantification of spikes in luminescence demonstrate that picrotoxin induces prolonged bouts of neuronal activity. n = 14-15 flies/condition. Two-group two-tailed t-test was used. *p<0.05. Data are presented as mean values ± SEM.

**Ext. Data Fig. 8.**
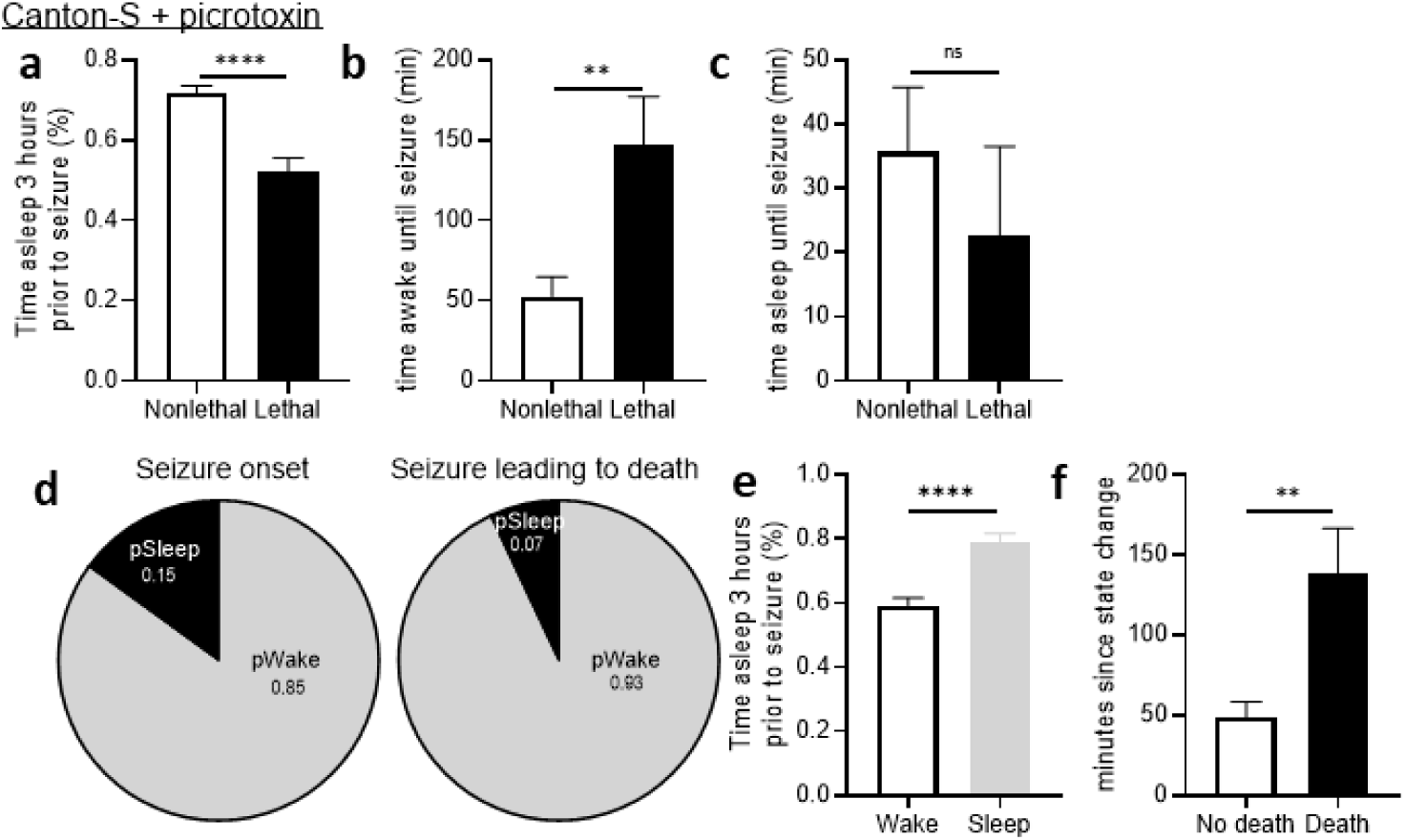
A new protocol for detection of spontaneous seizures in *Drosophila* reveals spontaneous seizures after sleep restriction. **a**, Retrospective analysis of wild-type Canton-S flies treated with picrotoxin demonstrates that lethal seizures occur after decreased sleep in the 3 preceding hours as compared to nonlethal seizures. n=41-44 seizures/condition. **b**, On average, wild-type Canton-S flies treated with picrotoxin with lethal seizures had longer wake times prior to seizure onset. n=34-38 seizure/condition. **c**, The duration of sleeping in wild-type Canton-S flies asleep at seizure onset did not predict seizure lethality. n=3-10 seizures/condition. **d**, Wild-type Canton-S flies treated with picrotoxin were more likely to have seizures during wakefulness, and lethal seizures were more likely to occur during wakefulness. **e**, Seizures occurring in sleep correlate with a greater percentage of the preceding 3 hours spent in sleep. n = 13-72 seizures/condition. **f**, Lethal seizures are more likely to occur as duration since a wake-to-sleep or sleep-to-wake transition increases. n = 41-44 seizures/condition. Two-group two-tailed t-test was used. **p<0.01, ****p<0.0001. Data are presented as mean values ± SEM.

**Ext. Data Fig. 9.**
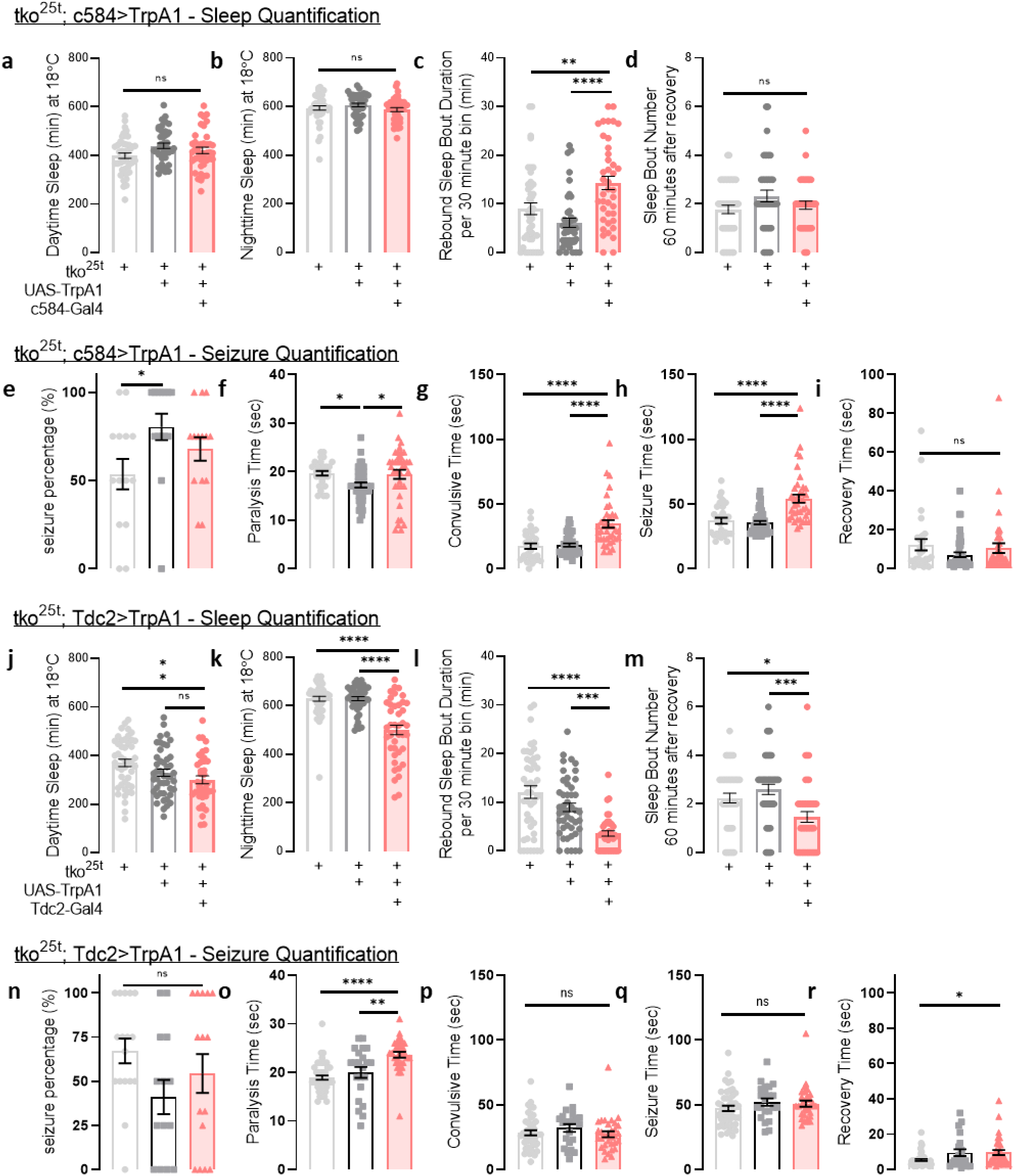
Sleep restriction using genetic manipulations leads to more severe induced seizures in *tko*^25t^ mutant flies. **a**, Before shifting to 30°C, there is no difference in daytime sleep in *tko*^25t^; c584-Gal4>UAS-TrpA1 flies. **b**, After shifting back from 30°C, there is no difference in nighttime sleep. **c**, After returning to 18°C, there is a prolongation of sleep bout duration in the ensuing 60 minutes, indicative of increased sleep consolidation. n = 40-43 flies/condition. **d**, After returning to 18°C, there is no change in the total number of sleep bouts in the ensuing 60 minutes. n = 40-43 flies/condition. **e-i**, *tko*^25t^; c584-Gal4>UAS-TrpA1 flies exhibit prolonged seizures after TrpA1 activation at 30°C for 24 hours. n = 14 vials/condition. n = 30-45 flies/condition. **j**, Before shifting to 30°C, daytime sleep in *tko*^25t^; Tdc2-Gal4>UAS- TrpA1 flies is comparable to genetic controls. **k**, After shifting back from 30°C, there continues to be a decrease in nighttime sleep in *tko*^25t^; Tdc2-Gal4>UAS-TrpA1 flies. **l**, After returning to 18°C, there is a decrease in sleep bout duration in the ensuing 60 minutes. n=41-46 flies/condition. **m**, After returning to 18°C, there is a decrease in the total number of sleep bouts in the ensuing 60 minutes. n=41-46 flies/condition. **n-r**, *tko*^25t^; Tdc2-Gal4>UAS-TrpA1 flies exhibit no change in convulsive (tonic/clonic) times and seizure times after TrpA1 activation at 30°C for 24 hours but show prolonged paralysis and recovery times. n = 14-17 vials/condition. n = 22-45 flies/condition. One-way ANOVA with Dunnett’s or Tukey’s multiple comparisons test or Kruskal-Wallis with Dunn’s multiple comparisons (for seizure percentage) test was used. *p<0.05, **p<0.01, ***p<0.001, ****p<0.0001. Data are presented as mean values ± SEM.

**Ext. Data Fig. 10.**
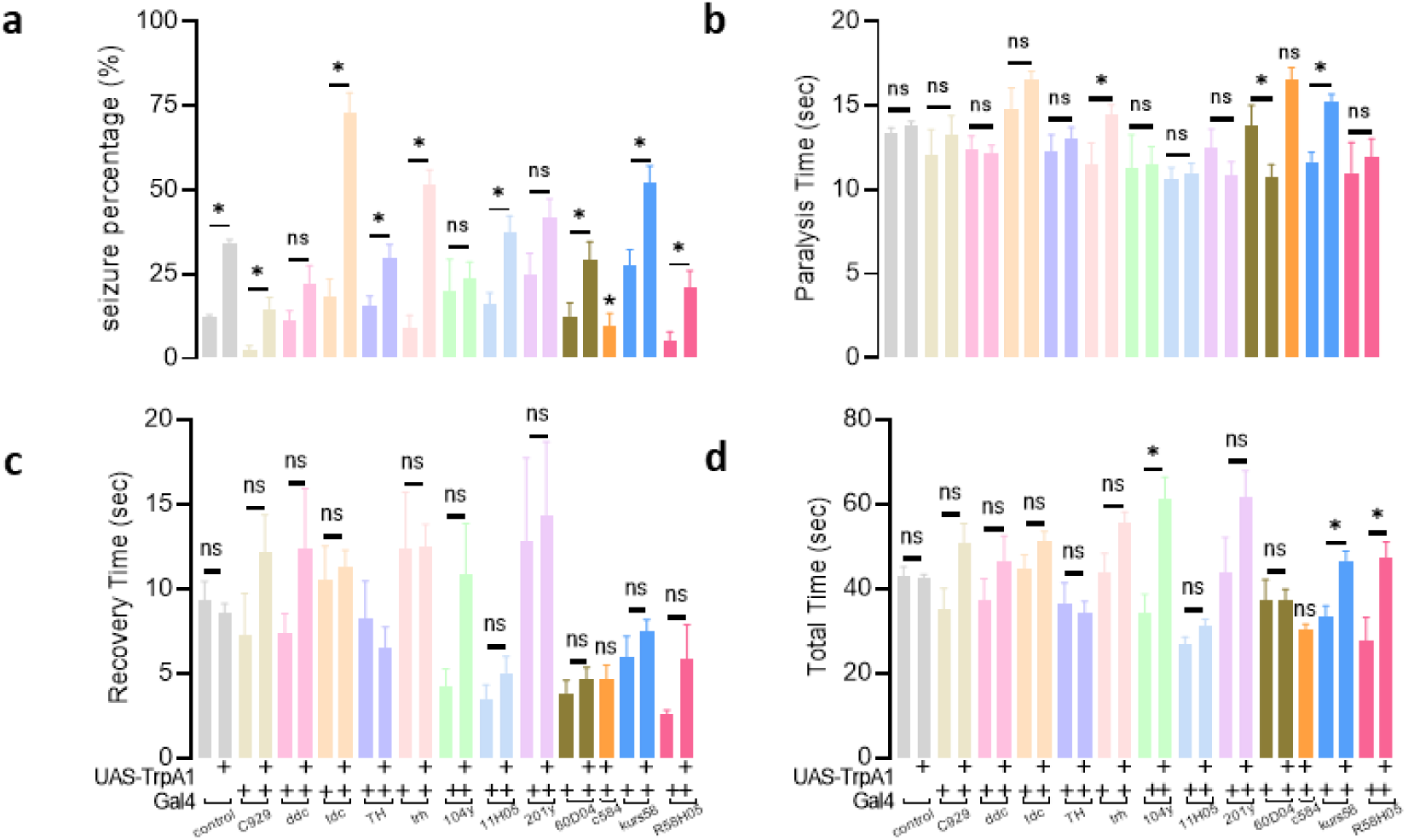
A Gal4 screen of wake- and sleep-promoting circuits and cellular subpopulations reveals that acute activation of sleep-promoting circuits worsens seizures. To drive TrpA1, Gal4 drivers were selected for peptidergic neurons (C929-Gal4), serotonergic/dopaminergic neurons (Ddc-Gal4), octopaminergic neurons (Tdc2-Gal4), dopaminergic neurons (TH-Gal4), serotonergic neurons (trh-Gal4), dorsal fan-shaped body (104y-Gal4), wake-promoting neurons (11H05-Gal4, 60D04-Gal4, c584-Gal4), mushroom body (201y-Gal4), pars intercerebralis (kurs58-Gal4), and ellipsoid body (R58H05-Gal4). **a**, Percentage of flies exhibiting seizures after mechanical stimulus. **b**, Paralysis time after seizure induction **c**, Recovery time after seizure induction. **d**, Total time until recovery after seizure induction. n = 4-77. Values shown are means and standard errors of the mean. Mann-Whitney test (for seizure percentages) or two-group two-tailed t-test was used. *p<0.05. Data are presented as mean values ± SEM.

**Ext. Data Fig. 11.**
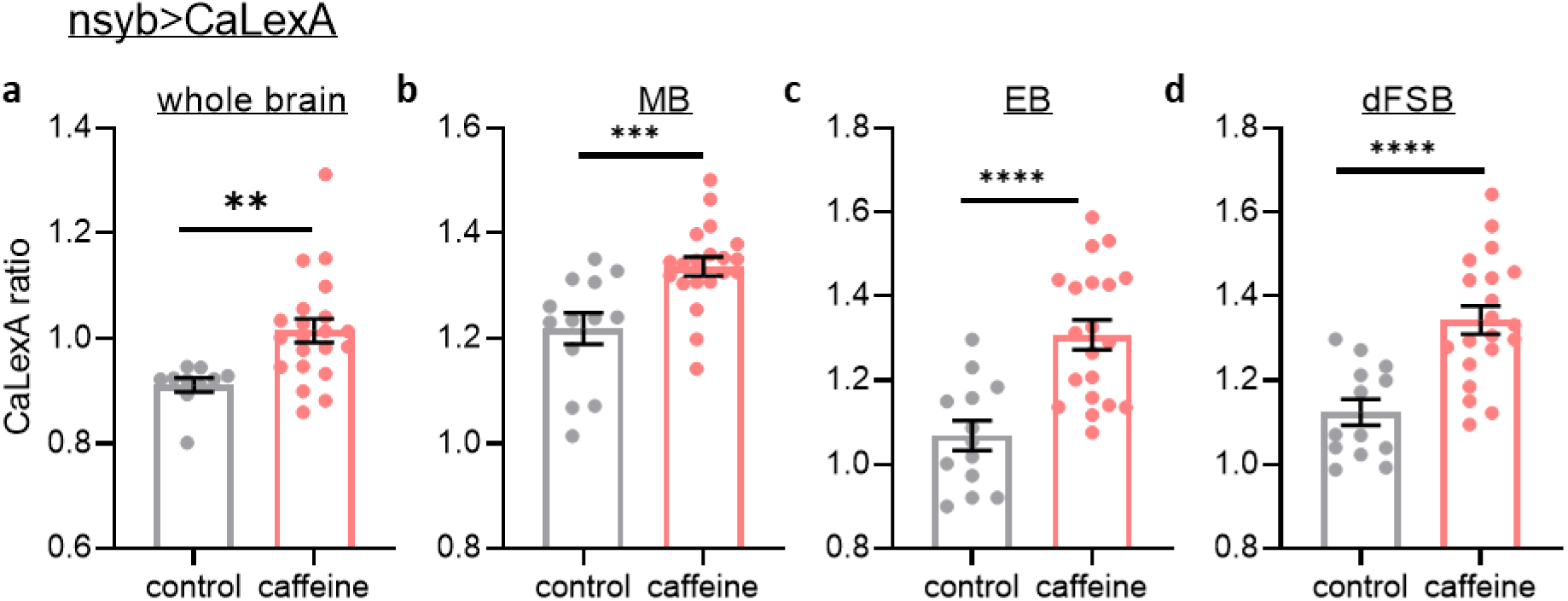
Activity is increased across whole brain, including sleep-promoting circuits, after sleep restriction. Brains dissected from nsyb-Gal4>CaLexA flies treated with vehicle or caffeine. **a-d**, Sleep restriction with caffeine in nsyb-Gal4>CaLexA flies leads to increased whole brain, mushroom body (MB), ellipsoid body (EB), and dorsal fan-shaped body (dFSB) CaLexA (GFP:RFP) signal. n=10-21 flies/condition. Two-group two-tailed t-test. **p<0.01, ***p<0.001, ****p<0.0001. Data are presented as mean values ± SEM.

**Ext. Data Fig. 12.**
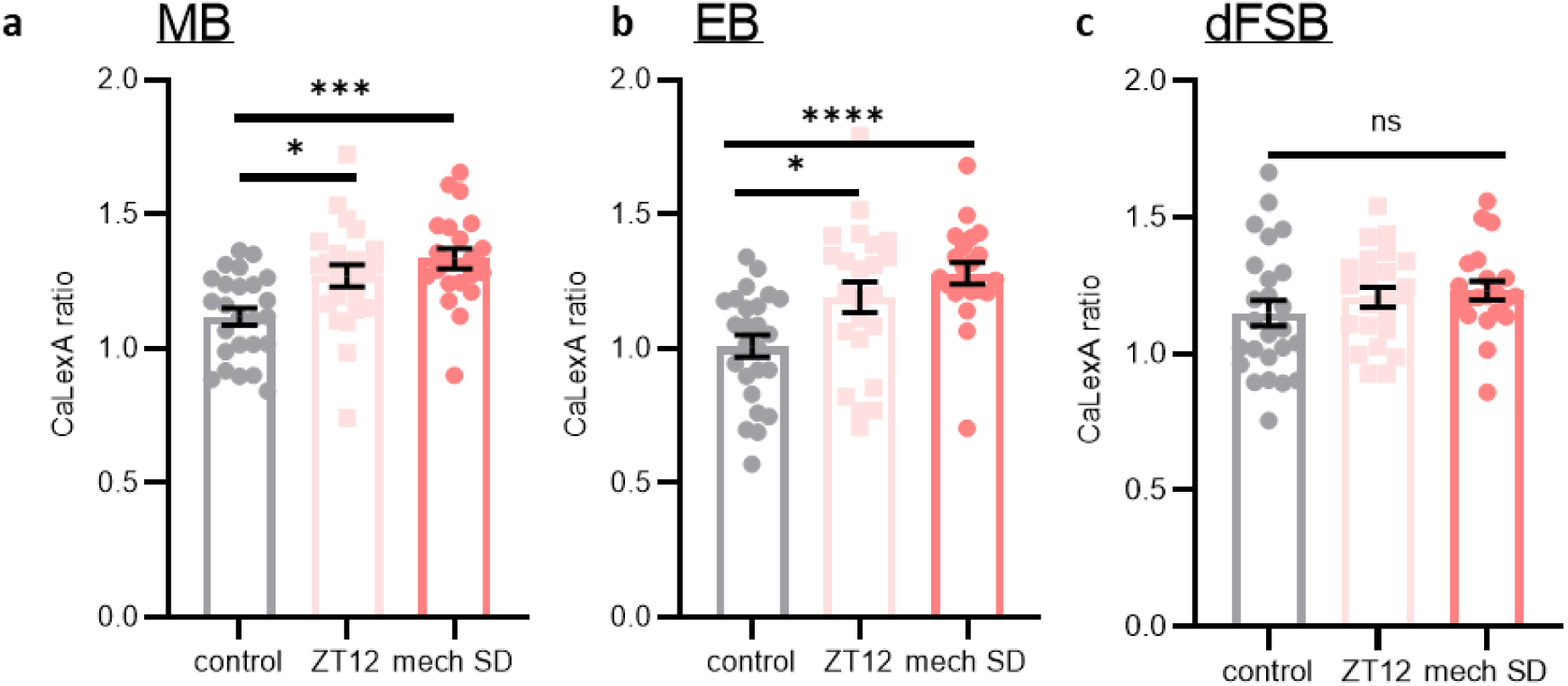
High sleep need correlates with increased activity of sleep-promoting brain regions. **a-c**, At the end of the day (ZT12) and after a night of mechanical sleep deprivation in nsyb-Gal4>CaLexA flies, there is increased CaLexA (GFP:RFP) signal in the mushroom body (MB) and ellipsoid body (EB) but not in the dorsal fan-shaped body (dFSB). n = 21-25 brains/condition. One-way ANOVA with Tukey’s multiple comparisons test was used. **p<0.01, ****p<0.0001. Data are presented as mean values ± SEM.

**Ext. Data Fig. 13.**
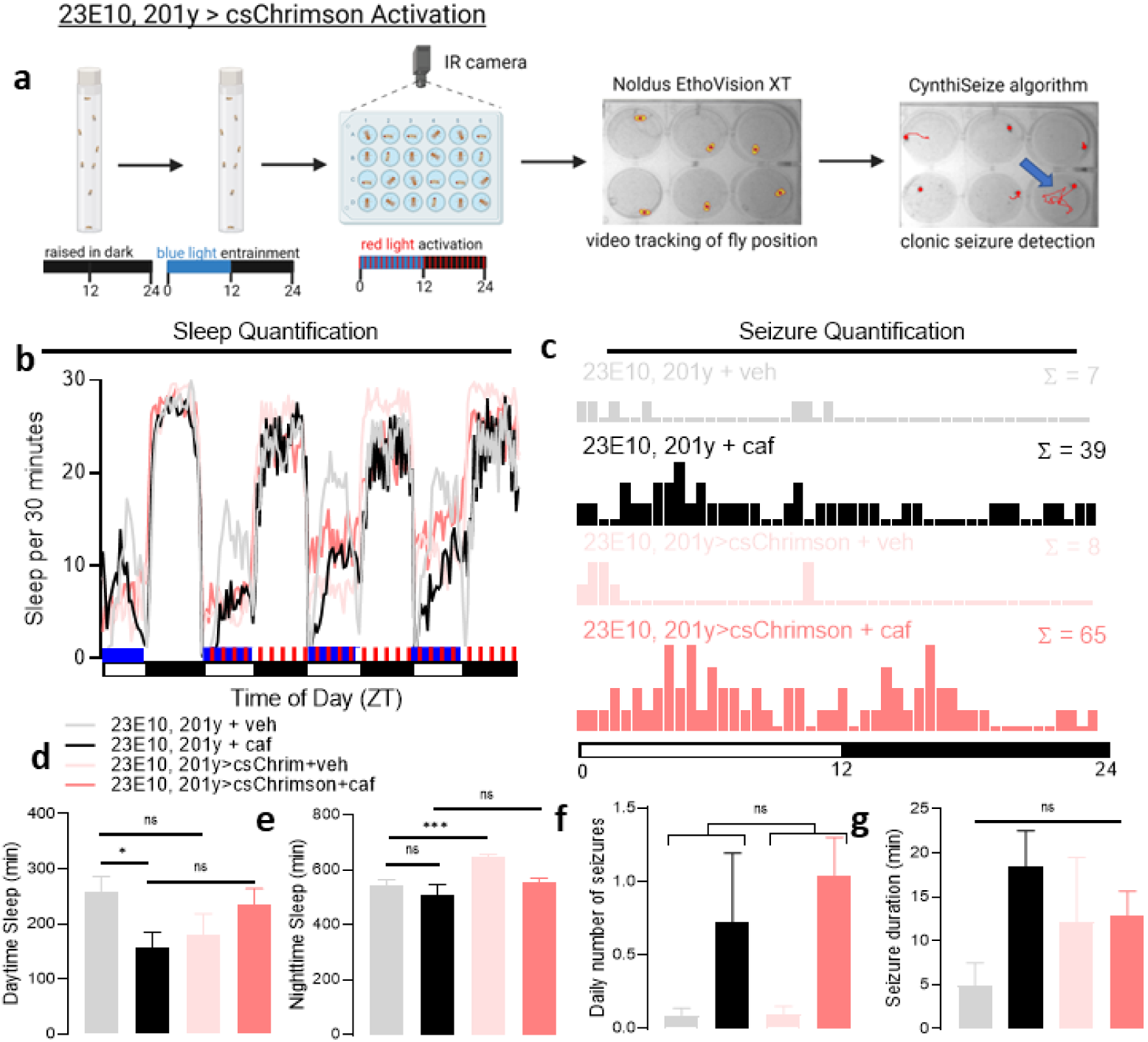
Optogenetic modulation of sleep-promoting circuits controls seizure risk in the setting of sleep loss. a, Experimental protocol showing flies were raised in darkness, entrained in blue light, then placed into 24- or 48-well plates for chronic video monitoring with red light stimulation. Fly positions over time were converted into XY coordinates, and a “CynthiSeize” algorithm was developed to identify seizures. **b**, **d**, **e**, Sleep quantification after 23E10-Gal4, 201y-Gal4>csChrimson activation. Caffeine decreases mean daytime sleep duration, and 23E10-Gal4, 201y-Gal4>csChrimson increases mean nighttime sleep duration. n=17-18 flies/condition. **c**, **f**, **g**, Seizure quantification after 23E10-Gal4, 201y-Gal4>csChrimson activation. Caffeine increases seizure frequency and is not additive with 23E10-Gal4, 201y- Gal4>csChrimson activation. n=17-18 flies/condition. One-way ANOVA with Dunnett’s T3 multiple comparisons adjustment, Kruskal-Wallis test with Dunn’s multiple comparisons adjustment, negative binomial model with Wald test (for daily number of seizures), or mixed effects model (for seizure durations) was used. *p<0.05, ***p<0.001. Data are presented as mean values ± SEM.

**Ext. Data Fig. 14.**
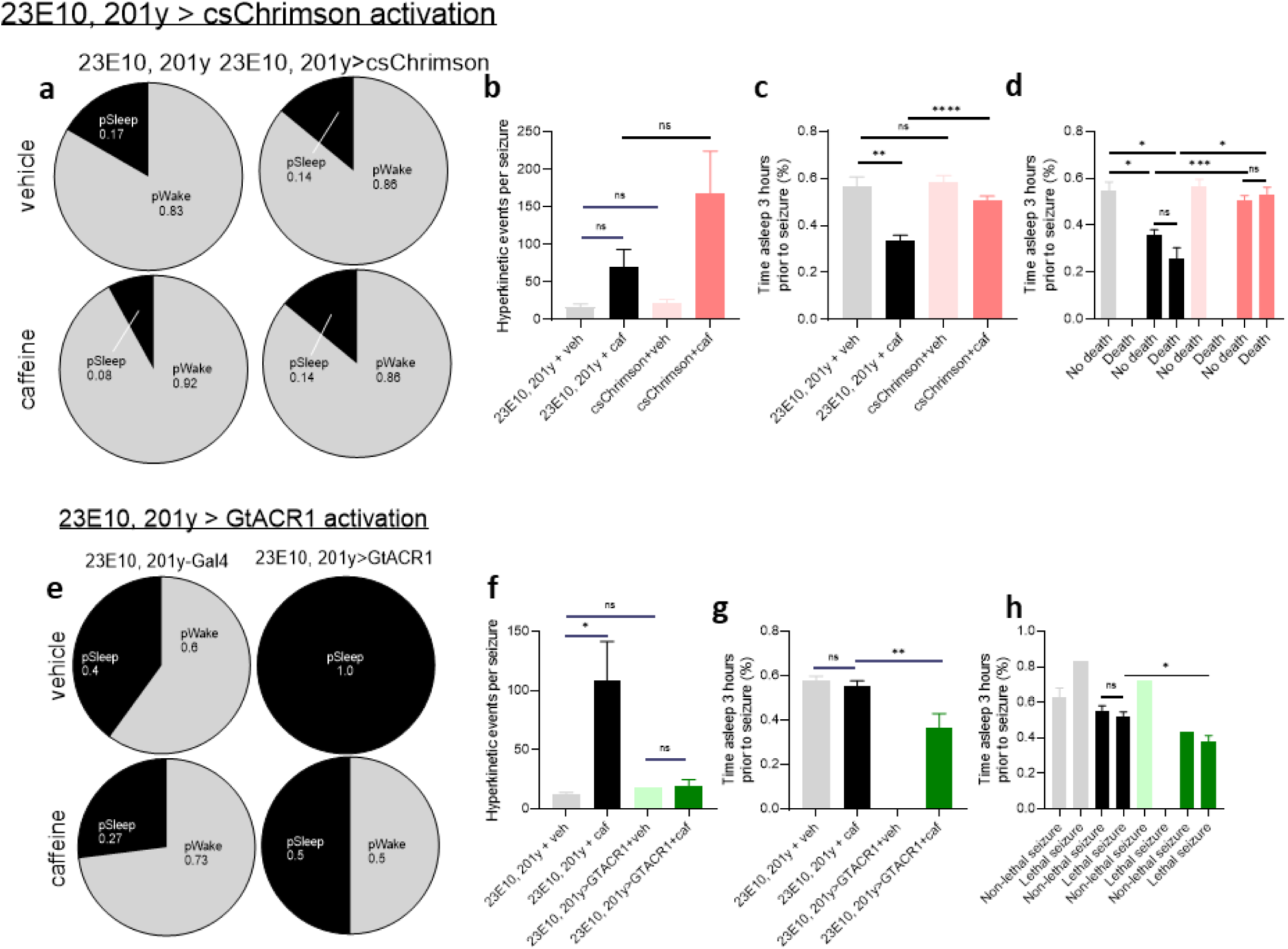
Optogenetic modulation of sleep-promoting circuits changes seizure risk. **a**, Likelihood of being awake (pWake) or asleep (pSleep) at seizure onset. Seizures are more likely to occur during wakefulness even after csChrimson activation or after caffeine treatment. **b**, There is no significant evidence that the number of hyperkinetic events per seizure change after caffeine treatment or after csChrimson activation. n = 6-49 seizures/condition. **c**, Seizures occurring with caffeine treatment during wakefulness are more likely to occur after decreased sleep in the 3 hours preceding seizures. Compared to 23E10-Gal4, 201y-Gal4 genetic control flies treated with caffeine, 23E10-Gal4, 201y-Gal4>csChrimson activation with caffeine treatment increases sleep quantity prior to seizure onset. n = 5-42 seizures/condition **d**, In flies treated with caffeine, lethal and non-lethal seizures are more likely to occur after decreased sleep in the preceding 3 hours. csChrimson activation increases sleep prior to seizure onset. n = 0-36 seizures/condition. **e**, Likelihood of being awake (pWake) or asleep (pSleep) at seizure onset. After 23E10-Gal4, 201y-Gal4>GtACR1 activation, a high proportion of seizures occur during sleep than without GtACR1 activation. **f**, Seizures occurring with caffeine treatment exhibit an increased number of hyperkinetic events per seizure n = 1-15 seizures/condition. **g**, In awake flies, GtACR1 activation with caffeine treatment decreases the sleep amount prior to seizure onset. n = 0-11 seizures/condition. **h**, Among flies treated with caffeine, flies with lethal seizures sleep less with GtACR1 activation. n = 0-8 seizures/condition. One-way ANOVA with Dunnett’s T3 multiple comparisons test or mixed effects model (for hyperkinetic events per seizure) was used. *p<0.05, **p<0.01, ***p<0.001, ****p<0.0001. Data are presented as mean values ± SEM.

**Ext. Data Fig. 15.**
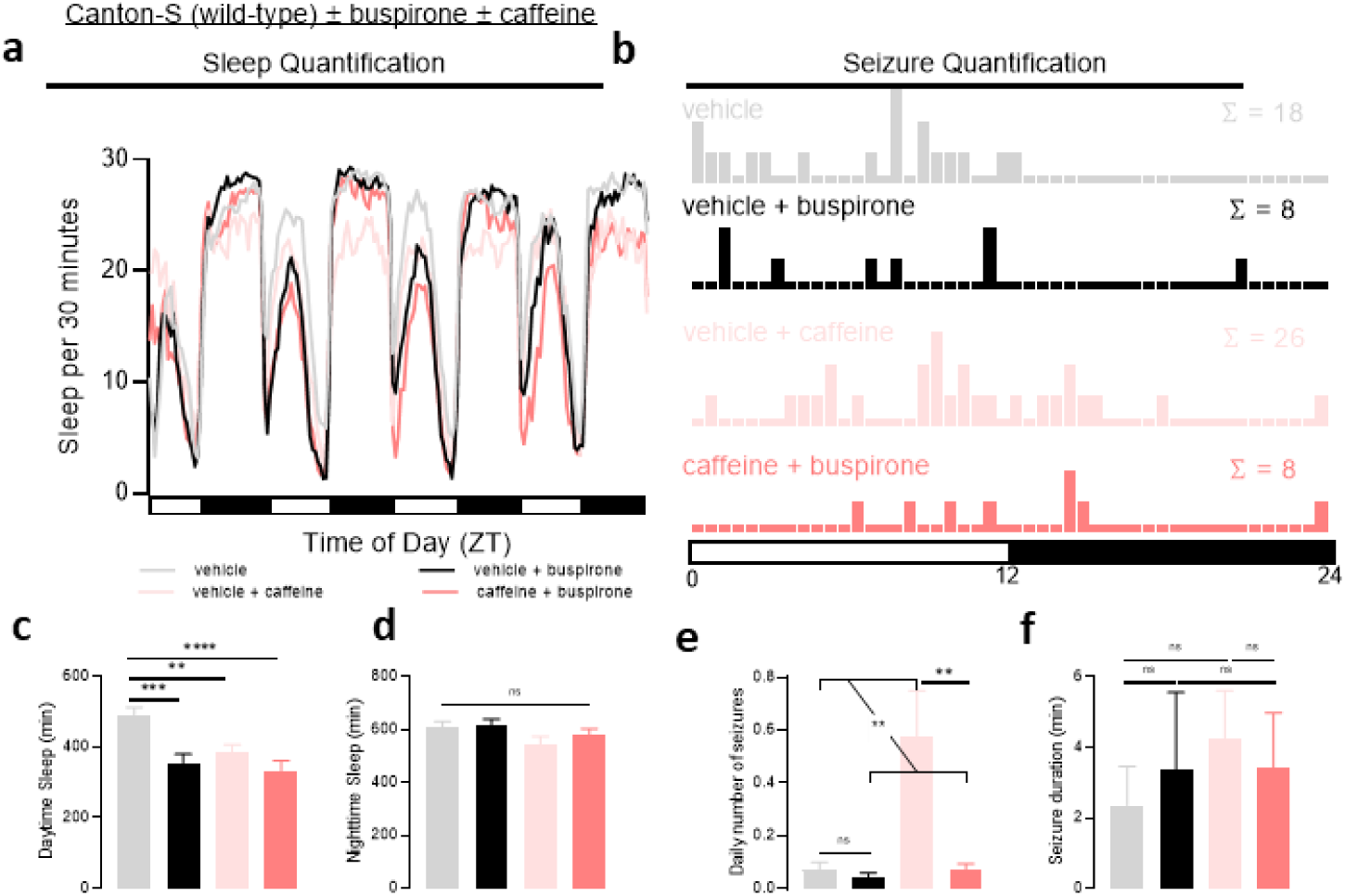
Increasing 5HT1A activity with buspirone can protect against seizures. **a, c, d**, Sleep quantification after administration of caffeine demonstrates decreased sleep, but no significant change with administration of buspirone, an FDA-approved 5HT1A agonist. n=36. **b, e, f**, Seizure severity is increased with caffeine, but reduced to baseline levels with co-administration of buspirone. n = 36. One-way ANOVA with Dunnett’s T3 multiple comparisons test, paired two-tailed t-test, negative binomial model with Wald test (for daily number of seizures), or mixed effects model (for seizure durations) was used. *p<0.05, **p<0.01, ***p<0.001, ****p<0.0001. Data are presented as mean values ± SEM.

**Ext. Data Fig. 16.**
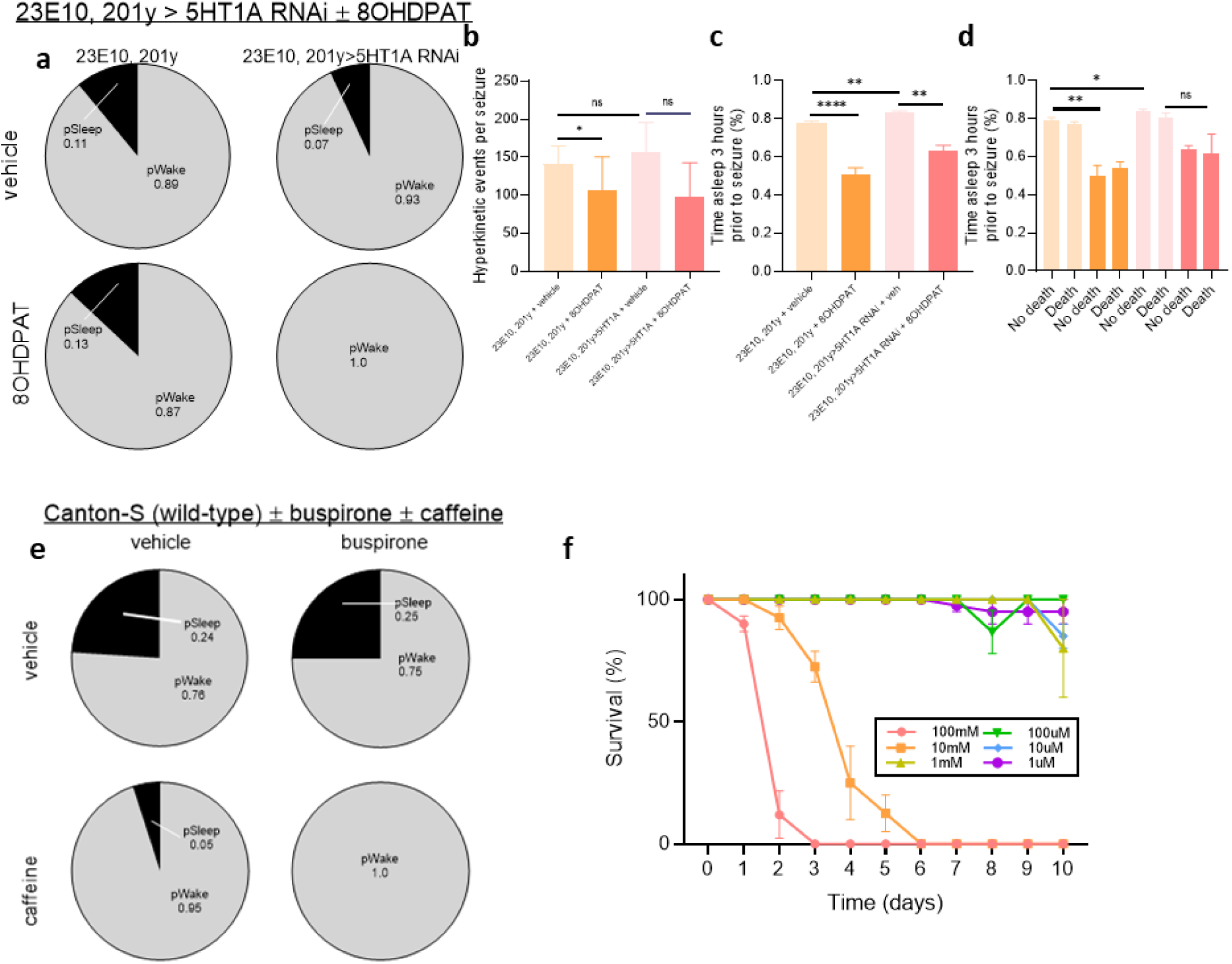
Serotonergic modulation of sleep-promoting circuits changes seizure risk. **a**, Likelihood of being awake (pWake) or asleep (pSleep) at seizure onset. Seizures are more likely to occur during wakefulness even after 8-OH-DPAT, a selective 5HT1A receptor agonist, treatment or 5HT1A RNAi-mediated knockdown. **b**, There is no significant evidence of a change in the number of hyperkinetic events per seizure after 8-OH-DPAT treatment or 5HT1A RNAi-mediated knockdown. n = 6-76 seizures/condition. **c**, 8-OH-DPAT treatment decreases the duration of sleep in awake flies prior to seizure onset. n = 6-68 seizures/condition. **d**, 8-OH-DPAT treatment decreases the duration of sleep prior to onset of lethal and non-lethal seizures. n = 2-46 seizures/condition. **e**, Likelihood of being awake (pWake) or asleep (pSleep) at seizure onset. Seizures are more likely to occur during wakefulness even after buspirone treatment. **f**, Wild-type Canton-S flies were treated with 1 uM, 10 uM, 100 uM, 1 mM, 10 mM, and 100 mM buspirone for 10 days and the number of surviving flies were counted. n = 4 vials/condition. One-way ANOVA with Dunnett’s T3 multiple comparisons test or mixed effects model (for hyperkinetic events per seizure) was used. *p<0.05, **p<0.01, ***p<0.001, ****p<0.0001. Data are presented as mean values ± SEM.

**Ext. Data Fig. 17.**
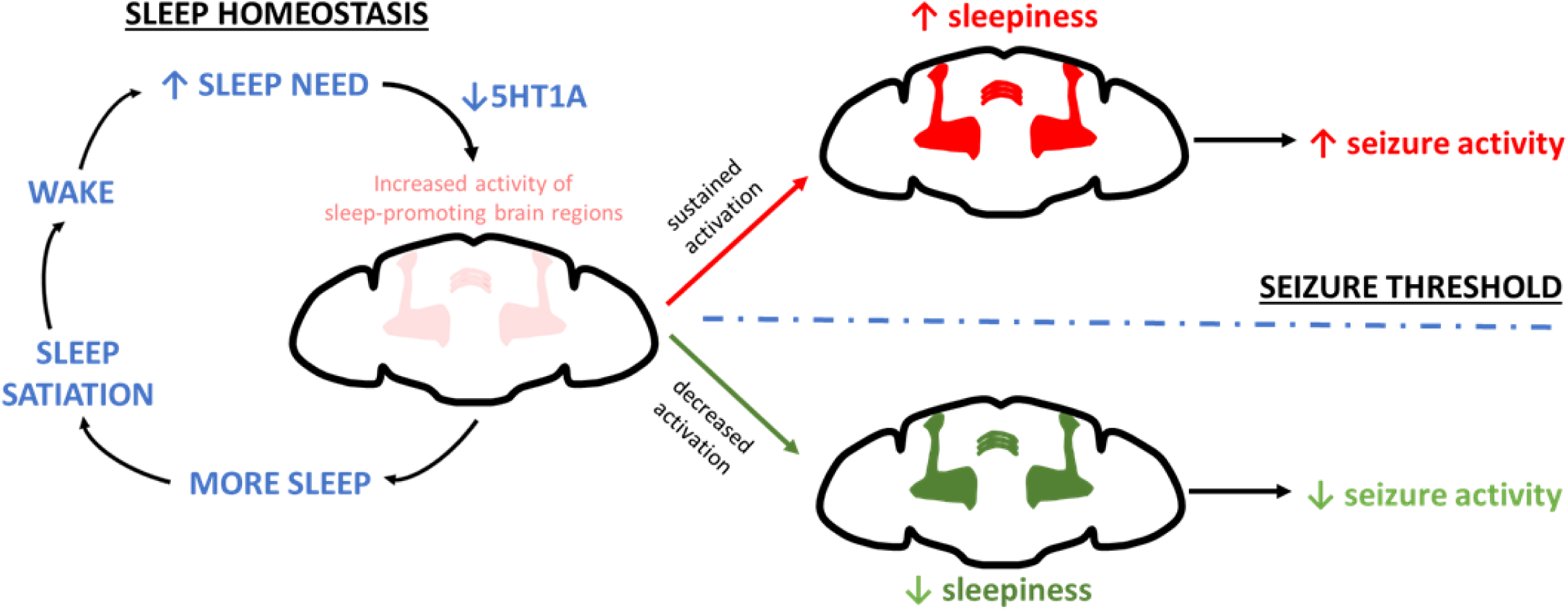
Schematic depicting how homeostatic sleep need drives increased seizure risk. Sleep need leads to increased activity of sleep-promoting circuits, and we find that this, in part, is driven by decreased expression of 5HT1A in the dorsal fan-shaped body (dFB). Increased activity of sleep-promoting circuits then leads to increased sleep drive, which allows flies to reach the required amount of sleep. This process is known as sleep homeostasis. A “price” of sleep homeostasis is that sustained activation sleep-promoting circuits leads to increased “sleepiness” and increased seizure activity. On the other hand, if the activity of sleep- promoting circuits is decreased (e.g. experimentally with GtACR1 activation or pharmacologically with 5HT1A agonism), seizures are less likely to occur.

