## Supplementary Information for "Sleepiness, not total sleep amount, increases seizure risk"

### Supplementary Methods

Drosophila melanogaster

Several existing mutant fly lines were used including two bang-sensitive mutants (*tko*$^{25t}$ and *eas*$^{pc80f}$), two short-sleeping mutants (*rye* and *sss*$^{P1}$), and *nemuri* mutants (*nur*$^{3}$). Wild-type Canton-S flies were used as unless otherwise indicated. UAS, Gal4, lexA, and lexAOp fly lines already not in lab or gifted were obtained from Bloomington Drosophila Stock Center (BDSC) in Indiana, USA or Vienna Drosophila Resource Center (VDRC) in Austria. See below for details on where each line was obtained from.

The following fly stocks were used in this study: *iso*$^{31}$ (control strain, lab stock), Canton-S (control strain, lab stock), *redeye* [1] (lab stock), *tko*$^{25t}$ [2] (gift from Dr. Dan Kuebler), *para*$^{bss1}$ (gift from Dr. Dan Kuebler), *eas*$^{pc80f}$ (lab stock), *sleepless* P1 [3] (lab stock), 60D04-Gal4 (BDSC #45356), nsyb-Gal4, (lab stock), 11H05-Gal4 (BDSC #45016), UAS-TrpA1 (II) [4] (gift from Dr. Leslie Griffith), R23E10-Gal4 (BDSC #49032), UAS-nGFP (BDSC #4775), w; UAS-CD8::RFP, LexAop-CD8::GFP-2A-CD8::GFP; UAS-mLexA-VP16-NFAT, LexAop-CD2::GFP [5, 6] (UAS-CaLexA; lab stock), c584-Gal4 (lab stock), Tdc2-Gal4 (lab stock), C929-Gal4 (lab stock), Ddc-Gal4 (lab stock), TH-Gal4 (BDSC #8848), trh-Gal4 (BDSC #38389), 104y-Gal4 (lab stock), 201y-Gal4 (lab stock), kurs58-Gal4 (BDSC # 80985), R58H05-Gal4 (BDSC # 39198), UAS-GCaMP7b (BDSC # 80907), 23E10-lexA (lab stock), LexAOp-csChrimson (lab stock), UAS-csChrimson (lab stock), UAS-GtACR1 (BDSC # 92983), UAS-5HT1A RNAi (VDRC #106094), *nur*$^{3}$ [7] (lab stock), UAS-nemuri [7] (lab stock), elav-GeneSwitch, (lab stock).

Induced seizure analysis

For induced seizures, bang-sensitive mutant flies were placed in groups of 4 flies into vials containing agar (2%) (Fisher NC1429200) and sucrose (5%) 24-hours prior to seizure induction. This was done because anesthetization of flies with $CO_2$ shortly before seizure induction caused flies to be refractory to seizures. Vials of all experimental conditions were struck on a countertop together four times. All vials were then vortexed on a Fisher Vortex Genie 2 (#12-812) at setting '5'. *tko*$^{25t}$ flies were vortexed for 5 seconds. *eas*$^{pc80f}$ flies were vortexed for 5 seconds. Vials were then placed in front of a USB camera with varifocal manual lens (Mokose #UC70-6-12MM) with video recording on a PC using OBS Studio 29.0. Videos were then subsequently analyzed for percentage flies with seizure in each vial, duration in atonic ("paralysis") phase, duration in tonic/clonic ("convulsive") phase, and duration in postictal ("recovery") phase for each individual fly. Scorers were blinded when possible.

For drug treatment experiments, prior to seizure induction, flies were treated with vehicle, caffeine (1 mg/ml for *tko*$^{25t}$ flies and 0.5 mg/ml for *eas*$^{pc80f}$ flies) [8], or gaboxadol (0.1 mg/ml) for 48 hours with drug mixed in agar (2%) and sucrose (5%). For *tko*$^{25t}$; elav-GeneSwitch>UAS-nemuri experiments, experimental flies with genetic controls were treated with RU486 (500 µM) for 72 hours prior to experiments.

For thermogenetic sleep deprivation, *tko*$^{25t}$; c584-Gal4>UAS-TrpA1 flies with genetic controls or *tko*$^{25t}$; Tdc2-Gal4>UAS-TrpA1 flies with genetic controls were raised at 18°C, maintained at 30°C for 24 hours, then allowed to equilibrate for 20 minutes at room temperature prior to mechano-sensitive seizure induction.

For TrpA1 [4] screening of sleep or wake promoting cells and circuits that affect seizure severity, experimental flies with genetic controls were raised at 18°C. Flies were then brought to

25°C for 4 minutes by immersing vials into a water bath. Vials were then quickly removed from the water bath for mechanical seizure induction, then returned to the water bath at 25°C with video recording of seizure percentages and durations.

Sleep quantification using multibeam infrared monitors

Flies were placed into locomotor tubes containing agar (2%) and sucrose (5%) with vehicle, caffeine (1 mg/ml for $tko^{25t}$ flies and 0.5 mg/ml $eas^{pc80f}$ flies), or gaboxadol (0.1 mg/ml).

Sleep was measured by counting the number of infrared beam breaks in *Drosophila* activity monitors, specifically the DAM5H multibeam monitor (Trikinetics). Locomotor data were collected using DAMsystem software (Trikinetics). Both 'Movement' and 'Counts' were assessed and extracted from raw files using DAMfilescan (Trikinetics). Sleep was defined as five minutes of inactivity [9, 10]. Sleep was then quantified using Insomniac 3.0 software [11].

For $tko^{25t}$; elav-GeneSwitch>UAS-nemuri experiments, experimental flies with genetic controls were treated with RU486 (500 µM) for 72 hours prior to sleep assessment. Locomotor tubes containing agar (2%) and sucrose (5%) with vehicle or RU486 (500 µM) were used.

Video tracking analyses of spontaneous seizures and sleep

Flies were placed into 24- or 48-well plates containing agar (2%) and sucrose (5%) with vehicle or drugs mixed into the agar. Caffeine was used at 1 mg/ml [8]. Picrotoxin dosing was determined with a toxicity assay (data not shown); for assays without caffeine, 0.5 mg/ml of picrotoxin was used, and for assays with caffeine, 0.05 mg/ml was used. 8-OH-DPAT, a selective 5HT1A receptor agonist, was used at 3 mM [12, 13]. Buspirone dosing was determined with a toxicity assay and used at 3 mM (Ext. Data Fig. 16f). Plates were then sealed with a plastic film (PerkinElmer TopSeal A Plus; 6050185), and small holes were placed through the film using a syringe to allow for air and humidity exchange. Plates were then placed in an incubator with light, temperature, and relative humidity control and illuminated with an infrared (IR) light. An IR camera (monochrome GigE camera with IR pass filter) was used for continuous monitoring through alternating 12-hour light:dark cycles at 25 frames per second for 96 hours per experiment.

EthoVision XT (Noldus Information Technology) was used to convert fly positions to XY coordinates over time. Thresholds for 'velocity', 'acceleration', 'mobility', 'mobility state', and 'movement' were developed in EthoVision XT to define a "hyperkinetic event" (HE). For bang-sensitive flies ($tko^{25t}$ and $eas^{pc80f}$), seizures were defined as 5 HE occurring in 50 seconds, with at least 7 HE per seizure. For picrotoxin-treated flies, seizures were defined as 5 HE occurring in 50 seconds, with at least 10 HE per seizure. In all cases, seizures that occurred within 15 minutes of each other were regarded as the same seizure. Threshold levels were set at values such that wild-type flies (Canton-S, $w^{1118}$, and $iso^{31}$) on vehicle (no caffeine or picrotoxin) never exhibited movements that were defined as seizures.

Movement was quantified as previously described [14]. Sleep was defined as inactivity for 5 minutes or more. Sleep or wake status was determined at the time of seizure occurrence. Time since state change is defined as time since a change from sleep-to-wake status or wake-to-sleep status. Sleep history is reported as the percentage of time asleep in the preceding 180 minutes.

For experiments involving optogenetic stimulation, experimental flies with genetic controls were raised in the dark. (1) For csChrimson experiments, flies were then placed in all-trans retinal (ATR) (300 µM) for 48 hours with 12-hour light:12-hour dark entrainment in blue light. After entrainment, flies were placed into 24- or 48-well plates containing picrotoxin, ATR, and caffeine with stimulation with red light for the first five minutes of every hour around the clock. (2) For GtACR1 experiments [15], flies were placed in all-trans retinal (ATR) (1 mM) for 48 hours with 12-hour light:12-hour dark entrainment in red light. After entrainment, flies were placed into 24- or 48-well plates containing picrotoxin, ATR, and caffeine with stimulation with green light for the first five minutes of every hour around the clock.

TRIC-luciferase assay

The nsyb-Gal4 driver was used to express UAS-TRIC-luciferase pan-neuronally. Adult flies aged 5-10 days were raised in a 12-hour light:12-hour dark cycle. Flies were fed 2 mM luciferin for 24 hours prior to the start of the experiment. Flies were then placed into a 96-well plate containing 100 uL of the following: agar (2%), sucrose (5%), luciferin (2 mM; Gold BioTechnology, Inc.), and picrotoxin (0.5 mg/mL). A clear adhesive plastic film (Top-Seal-A; Perkin Elmer) was used to cover the 96-well plate. Plates were then loaded into a Cytation 5 multimode reader (BioTek) with imaging of luminescence occurring through the thin agar layer and imaging of fly posture through the clear adhesive plastic film. Luminescence was detected with PMTs every 14 minutes. Bright field images were acquired with a 4x objective.

CaLexA imaging

For drug treatment, flies were treated with vehicle or caffeine (1 mg/ml) mixed into agar (2%) and sucrose (5%) [8] for 48 hours. For mechanical sleep restriction, flies were shaken on a pre-programmed vortexer for 12 hours overnight. Adult fly brains were dissected in cold phosphate buffered saline (PBS) and then fixed in 2% paraformaldehyde for 40 minutes at room temperature. Brains were then washed twice for 20 minutes in PBS with 0.3% Triton-X (PBST). Samples were then placed in 50% glycerol and mounted in Vectashield: H1000. Both control and experimental brain were mounted on the same slide for visualization. Primary GFP and RFP signal was visualized without signal amplification with antibodies. On the same day as mounting, brains were visualized on a Leica Stellaris 8 confocal microscope. Identical settings were used for laser intensity and gain for control and experimental conditions. After image acquisition, Fiji was used for image processing and analysis. All data are presented as GFP:RFP ratios. Whole brain calcium levels are presented as total signal from z-projections. Subregion analyses were performed as measurements of ROIs in individual slices.

RNA sequencing of the dorsal fan shaped body

To isolate and sort dFB neurons, we first generated flies with nuclear GFP expression using the R23E10-Gal4 (BDSC #49032) driver crossed with UAS-nGFP (BDSC#4775). Male flies aged 5-7 days were subjected to sleep deprivation overnight or regular nighttime sleep in the same incubator, and brains were dissected the next day at ZT0. The brains were dissociated using a protocol from Hongjie Li et al. 2018; briefly, flies were dissected in Schneider's medium, followed by dissociation in Papain solution and filtration through a 100 µm cell strainer. The ventral nerve cords were not included. The resulting cells were then suspended in Schneider's medium, and 100 GFP+ cells from each condition were sorted using either BD FACSMelody or BD FACSAria (BD Biosciences). Dead cells were excluded using 4', 6-diamidino-2-phenylindole (DAPI). Doublets were also excluded based on forward scatter (FSC-H by FSC-W) and side scatters (SSC-H by SSC-W). FSC-A determined the size of cells, and validation was done using cells from flies expressing nSyb>nGFP. The sorted cells were frozen and placed into a 96-well plate with lysis buffer from the Smart-seq2 HT kit.

The sorted cells were sent to Admera Health (admerahealth.com) for RNA extraction, RNA library construction, and sequencing using the Smart-seq2 HT kit. The sequencing data were then mapped to the fly genome (BDSG6) using Hisat2 (daehwankimlab.github.io/hisat2), and the alignment results were counted by LiBiNorm tool (warwick.ac.uk/fac/sci/lifesci/research/libinorm) based on the reference genome from GENCODE. Both raw count and TPM (transcripts per million) data were used separately in further analysis. The raw count data were analyzed by IDEP v0.95 (bioinformatics.sdstate.edu/idep) for genes expressed differentially. Genes with CPM > 0.5 were detected in at least three independent samples, and missing values treated as gene median was selected to filter out low-expressed genes. The regularized log transformation was applied to remove the dependence of the variance on the mean. The transformed raw count data were then used for further clustering and PCA. Differentially expressed genes were identified using DESeq2 with an FDR cutoff of 0.1 and minimum fold change of 2.

Quantification and statistical analysis

Pre-testing with Shapiro-Wilk test and Kolmogorov-Smirnov tests were conducted to assess normality and the choice of parametric or non-parametric testing. For two groups, unpaired or paired two-tailed t-test was used for data that were reasonably assumed to be approximately normally distributed. For two groups, if the variance was significantly different, unpaired two-tailed t-test with Welch's correction was used. When comparing two groups, a Mann-Whitney test was used if the normality assumption was not justified, including for ordinal data. When comparing a single control group to multiple experimental groups, a one-way ANOVA with Dunnett's multiple comparisons test was used for approximately normally distributed data, and a Kruskal-Wallis with Dunn's multiple comparisons test was used if the normality assumption was not justified, including for ordinal data. When comparing three or more groups with multiple comparisons, a one-way ANOVA with Tukey's multiple comparisons test was used for normally distributed data, and a Kruskal-Wallis with Dunn's multiple comparisons test was used when the normality assumption was not justified. If the variance was significantly different between groups when comparing three or more groups using Bartlett's test, one-way ANOVA with Dunnett's T3 multiple comparisons test was used with individual variances computed for each comparison. The number of flies awake or asleep when seizures occur is reported as a percentage with 'pWake' representing the number of flies awake at seizure onset/total number of seizures, and 'pSleep' representing the number of flies asleep at seizure onset/total number of seizures.

For hypothesis testing of spontaneous seizure frequency, given the categorical variable (seizure count per day per fly), right-skewed distribution of this dataset, and censoring due to fly death during the experiment, a negative binomial model with Wald test was implemented. Exploratory analyses implementing negative binomial models were run (1) to assess the effects of 8-OH-DPAT on seizure frequency for each genotype (Fig. 5f) and (2) to assess the effects of buspirone on seizure frequency with and without caffeine (Ext. Data Fig. 15e). For hypothesis testing of seizure duration and hyperkinetic events per seizure, in datasets where one fly experienced multiple seizures, a mixed-effects model was implemented to account for both intra- and inter-fly variability. To reduce the skewness of the original data, p-values were calculated using log-transformed seizure duration and hyperkinetic event count.

Mean and standard error of the mean are used to visually represent the data. Additional details about the sample size (n) for each experiment, statistical testing, and p-values are provided in the Figure Legends. GraphPad Prism was used for all statistical analyses, except for significance testing of spontaneous seizure frequency, seizure duration, and hyperkinetic events per seizure, which were performed in R version 4.2.1.

**Supplementary Discussion**

In *Drosophila*, across the 3 neurogenetic and 1 pharmacological models tested, most seizures occurred during wakefulness (Ext. Data Fig. 6b, f, Ext. Data Fig. 8d, Ext. Data Fig. 14a, Ext. Data Fig 16a, e). We find that the wake state correlates with a history of increased wakefulness (Ext. Data Fig. 8d, e), so whether the wake state or a history of wakefulness drives seizures during wakefulness in *Drosophila* remains unclear. The occurrence of seizures during wakefulness or sleep is also likely tied to etiology, e.g., nocturnal seizures are associated with sleep-related hypermotor epilepsy due to pathogenic variants in *CHRNA4* [16, 17]. Our findings also indicate that seizures occurring after recent transitions from wake → sleep or sleep → wake are more likely to be non-lethal (Ext. Data Fig. 8f). This is consistent with the finding that flies that have been awake for longer periods of time are more likely to have severe lethal seizures (Ext. Data Fig. 8b).

Our transcriptomic analysis of the dFB after sleep restriction revealed downregulation of 5HT1A. *Drosophila* express five serotonin receptors, including 5HT1A, 5HT1B, 5HT2A, 5HT2B, and 5HT7 [18]. The relationship between sleep and serotonin is complex; serotonin binds to multiple receptors in multiple cell types across the brain to produce varied effects on sleep [19]. When 5HT1A is globally disrupted in 5HT1A mutant flies, sleep becomes decreased and fragmented [20]. We show here that a selective 5HT1A agonist, 8-OH-DPAT, also decreases sleep, and that this is predominantly mediated by action at sleep-promoting cells (Fig. 5e); this further emphasizes the complex relationships between sleep and serotonin and the context dependence of the specific cell types and receptors involved in serotonergic signaling. These data are consistent with a model in which sleep restriction leads to loss of 5HT1A in sleep promoting centers. Given the known inhibitory effects of 5HT1A [13], loss of 5HT1A-medited inhibition would promote activity of sleep-promoting centers to lead to increased sleep. Consistent with this model, we demonstrate that RNAi-mediated knockdown of 5HT1A in sleep-promoting centers increases sleep (Fig. 5). Our manipulations of 5HT1A used the 23E10-Gal4 driver, which targets not only the dFB but also specific ventral nerve cord (VNC) cells that were recently shown to be critical for sleep effects [21, 22]. Given that we observed changes in 5HT1A in the dFB, we propose that the decrease in 5HT1A acts in conjunction with other sleep loss-induced changes in the dFB—decreased $K^+$ leak conductance [23] and increased Rho-GTPase-activating protein crossveinless-c activity [24]— to promote homeostatic sleep rebound. We also find that promoting 5HT1A activity in the sleep-promoting centers not only decreases sleep, presumably due to decreased activity of sleep-promoting centers, but also decreases seizure burden. Therefore, while increased activity of sleep-promoting centers is important for sleep homeostasis, it comes at the cost of promoting seizures if the neural milieu is seizure-prone (Ext. Data Fig. 17).
